# Kidney single-cell atlas reveals myeloid heterogeneity in progression and regression of kidney disease

**DOI:** 10.1101/2020.05.14.095166

**Authors:** Bryan R Conway, Eoin D O’Sullivan, Carolynn Cairns, James O’Sullivan, Daniel J. Simpson, Angela Salzano, Katie Connor, Peng Ding, Duncan Humphries, Kevin Stewart, Oliver Teenan, Neil C Henderson, Cecile Benezech, Prakash Ramachandran, David Ferenbach, Jeremy Hughes, Tamir Chandra, Laura Denby

## Abstract

The kidney has a limited capacity to repair following injury, however, the endogenous reparative pathways are not well understood. Here we employ integrated droplet- and plate-based scRNA-seq in the murine reversible unilateral ureteric obstruction model to dissect the transcriptomic landscape at the single cell level during renal injury and resolution of fibrosis. We generate a comprehensive catalogue of the changes induced during injury and repair, revealing significant myeloid cell heterogeneity, which would not have been identifiable by conventional flow cytometry. We identify new markers for the myeloid populations within the kidney as well as identification of novel subsets including an *Arg1*^+^ monocyte population specific to early injury and a *Mmp12*^+^ macrophage subset exclusive to repair. Finally, using paired blood exchange to track circulating immune cells, we confirm that monocytes are recruited to the kidney early after injury and are the source of *Ccr2*^+^ macrophages that accumulate in late injury. Our data demonstrate the utility of complementary technologies to identify novel myeloid subtypes that may represent therapeutic targets to inhibit progression or promote regression of kidney disease.

## Introduction

Chronic kidney disease (CKD) affects approximately 10% of the global population (1) and is a major risk factor for end-stage kidney disease and cardiovascular disease (2–4). It is characterised by the progressive loss of renal filtrative capacity, with functional renal tissue replaced by excess extracellular matrix (ECM). Indeed, tubulointerstitial fibrosis remains the best histological predictor of renal outcome, even in diseases considered predominantly glomerular (5).

It is now recognised that CKD is not always progressive, but that regression of albuminuria and improvement in renal function can occur if the stimulus provoking injury is removed such as by improving blood glucose or blood pressure control in patients with diabetes or hypertension (6–8). Furthermore, regression of established fibrosis has been observed following prolonged normalisation of blood glucose levels after successful pancreas transplantation (9, 10). Hence, kidney disease may progress or regress, depending on whether the chronicity or severity of the injury stimulus exceeds the endogenous reparative capacity of the kidney. However, the cellular and molecular pathways that mediate regression of the injury are not well understood, at least in part because renal biopsies are rarely performed in patients who are clinically improving.

The innate immune system has been implicated in both progression and regression of fibrosis in multiple organs including the kidney (11–15). Recruitment of pro-inflammatory monocytes (16, 17) to the injured kidney via CCL1-CCR2 signalling (18, 19) may exacerbate tissue damage through the release of pro-inflammatory factors and by activating myofibroblasts. Macrophages can also be injurious, but in addition they may mediate repair by scavenging cell debris, degrading excess extracellular matrix and by secreting factors that may promote regeneration of injured tissue (20–22). Tissue macrophages are heterogeneous and inherently plastic and may adopt different phenotypes in response to environmental cues. Most studies have used panels of cell surface markers to characterise myeloid cell subsets on flow cytometry; however, this approach is inherently biased and is unlikely to capture the full phenotypic spectrum. Recent advances in transcriptomics including single-cell RNA sequencing (scRNAseq) have been used to characterise the myeloid cells in the healthy, injured and regenerating kidney (23–25) and other organs (26–30). However, while macrophages may mediate regression of fibrosis (31), the macrophage heterogeneity during regression of fibrosis in the kidney remains uncertain.

In order to dissect the molecular pathways mediating injury and repair in the kidney, we have developed rodent models in which sustained renal injury induces renal fibrosis, but where the injury can be withdrawn to facilitate repair (32, 33). In Cyp1a1mRen2 rats, chronic hyperglycaemia and hypertension induces pro-fibrotic and immune gene expression, whereas subsequent tight glycaemic and blood pressure control down-regulates pro-fibrotic pathways while innate immune genes remain persistently elevated, with evidence of a switch towards a more reparative macrophage phenotype (32, 34). In the reversible unilateral ureteric obstruction (R-UUO) model, we (33) and others (35) have demonstrated regression of established tubulointerstitial fibrosis following reversal of obstruction, with partial persistence of interstitial macrophages. However, the macrophage heterogeneity during induction and regression of fibrosis is not yet established.

Here we utilise the R-UUO model to characterise the myeloid cell phenotypes observed in renal injury and repair. scRNAseq analysis of renal cortex identified myeloid cell subsets that were indistinguishable by standard flow cytometry markers, with the relative proportions of the subsets changing dynamically during injury and repair. Similar myeloid cell phenotypes are observed in the human kidney, suggesting that they may represent specific targets to slow progression of CKD or promote renal repair.

## Methods

### Animal models

All protocols and surgical procedures were approved by the Animal Ethics Committee, University of Edinburgh. Animal experiments were conducted in accordance with the Animals Scientific Procedures Act UK 1986, under Home Office project licenses 70/8093 and 70/8867.

### Reversible unilateral ureteric obstruction model (R-UUO)

The R-UUO model was performed as previously described (33). Briefly, 8 week old male C57BL/6JOlaHsd mice (Enviago) underwent laparotomy and the left ureter was isolated and the distal portion was ligated twice with 6/O black braided silk suture close to the bladder. In mice that required reversible ureteric obstruction, a silastic tube was placed around the ureter immediately proximal to the ligature to prevent excessive dilatation. Following 7 days of obstruction, the ureter was re-anastomosed into the bladder and the peritoneum and skin sutured closed. Mice were culled at day 2 or day 7 post-UUO or 7, 14 or 28 days following ureteric re-anastomosis after 7 days of obstruction by CO2 narcosis and dislocation of the neck. SMART-Seq2 studies utilised Macgreen mice (36), in which the Csf1r promoter drives enhanced green fluorescent protein as a reporter specific to myeloid cells.

### Paired Blood Exchange (PBE)

Male C57BL/6NCrl mice which are homozygous for CD45.2 were paired with male Ly5.1 mice (Charles River), which are CD45.1/CD45.2 heterozygotes. 4 pairs of mice were used. The PBE was performed as previously described (37). Briefly, all animals had a right jugular venous catheter inserted prior to UUO surgery. 1 day post-UUO, 15 x 150μL aliquots of whole blood were exchanged between each animal in the pair over a 20 minute period. Two pairs were culled at both 2 days and 7 days post-UUO and whole blood and kidney tissue (UUO and contralateral kidney) was harvested.

### Immunohistochemistry

Kidney tissue was fixed and FFPE 4μm tissue sections prepared. Sections were rehydrated and staining performed using the Sequenza system (Thermo Scientific). Sections were incubated with avidin/biotin blocking kit (SP2001, Vector Labs) and blocked with serum-free protein block (X0909, Dako). Tissue sections were incubated with primary antibody (Suppl. Table 1), diluted in antibody diluent (S202230, Dako UK Ltd), overnight at 4°C before incubation with biotinylated secondary antibody (Suppl Table 1) for 30 minutes at room temperature. Vectastain RTU ABC reagent (PK7100, Vector Labs), was then applied followed by incubation with DAB + substrate chromogen system (K3468, Dako) and then counterstained with hematoxylin before dehydration and mounting with Pertex mounting media (3808707E, Histolab Products AB). The stained section was scanned with a Zeiss Axio Scan.Z1 Slide Scanner (Carl Zeiss Microscopy). The % of DAB staining per section (n=6-8/gp) was determined by ImageJ.

### Immunofluorescence

Slides were de-waxed in xylene (2×5min) and rehydrated and antigen retrieved (7 minutes at 60% power microwave). Slides were allowed to cool at room temperature, mounted in Sequenza (ThermoFisher) racks, rinsed twice in phosphate-buffered saline (PBS) and blocked for 45 minutes in Gentex block (120μl) at room temperature. Primary antibodies were incubated at the concentrations in Suppl Table 1 in antibody diluent (Abcam) and incubated overnight at 4°C. Slides were then washed twice with PBS and secondary antibodies added at a concentration of 1:200 diluted in antibody diluent and incubated at room temperature for 30 minutes. Slides were again washed twice with PBS and then blocked with Gentex block for 45 minutes at room temperature. The second primary antibody was added to the slides and they were incubated at 4°C overnight. Slides were washed and secondary antibodies applied. For dual immunofluorescence, the slides were boiled in 10mM citrate and blocked with serum-free protein block for 1hr prior to incubation fo the second primary antibody. The antibodies were visualized by incubation with Tyramide Red or Green (Perkin Elmer) for 10 minutes. After washing slides were mounted with DAPI Fluoromount-G^®^ (Southern Biotech) and a coverslip applied prior to visualisation.

### Bone-marrow derived macrophage (BMDM) culture

C57BL/6JOlaHsd mice hind legs were removed before skin and underlying muscle excised with sterile scissors and forceps to isolate the femur. The bone marrow was then flushed out in DMEM (Gibco) containing 10% L929 conditioned media, 10% fetal calf serum and 1% Penicillin-Streptomycin. Cell suspensions were cultured for 1 week in 60mL sterile Teflon pots at 37°C, 5% CO_2_. Macrophages were then plated into 6 well plates and incubated with 10μg FITC conjugated collagen (D12052) and left overnight. Cells were collected and run through flow cytometry to quantify the FITC signal.

### RNA extraction, gene expression and bulk RNA sequencing

Total RNA from cortical kidney tissue was isolated using the RNeasy kit (Qiagen, Hilden, Germany) following the manufacturer’s instructions. For qPCR analysis of targeted gene expression, cDNA was synthesised from 1μg of template RNA using the QuantiTect Reverse Transcription Kit (QIAGEN, Venlo, Netherlands). qPCR was performed using the PerfeCTa FastMix II probemaster (VWR, UK) and TaqMan Gene Expression assay specific primers (Life Technologies, Suppl. Table 2) and normalised to hypoxanthine-guanine phosphoribosyltransferase (*Hprt1*).

Prior to RNA sequencing, RNA integrity was checked using Agilent Nanochips, with only samples with RIN >7 utilized in subsequent analysis. 4 mice/group underwent RNAseq, with the animals selected on the basis that their *Havcr1* gene expression as determined by qRT-PCR was closest to the mean of that group. A PolyA library was constructed and run on a HiSeq2500 using 2×100bp paired-end sequencing. FastQC was used for initial quality control, reads were mapped to the mm10 transcriptome using RSEM and Bowtie2, while DESeq2 was used for differential gene expression analysis. Data was deposited in the National Center for Biotechnology Information Gene Expression Omnibus database (accession #GSE145053). The shinyNGS R package was used to generate gene clusters in Fig. 1c. Genes with an average FPKM of <1 across all groups were excluded from analysis. Genes which were not significantly differentially expressed (adjusted p-value <0.05, determined using DESeq2 R package) in any group (compared to sham) were excluded from analysis. Using the feature-based clustering module of the shinyNGS package, 7810 genes were assigned to one of six clusters based on expression change between each of the groups.

**Fig. 1:**
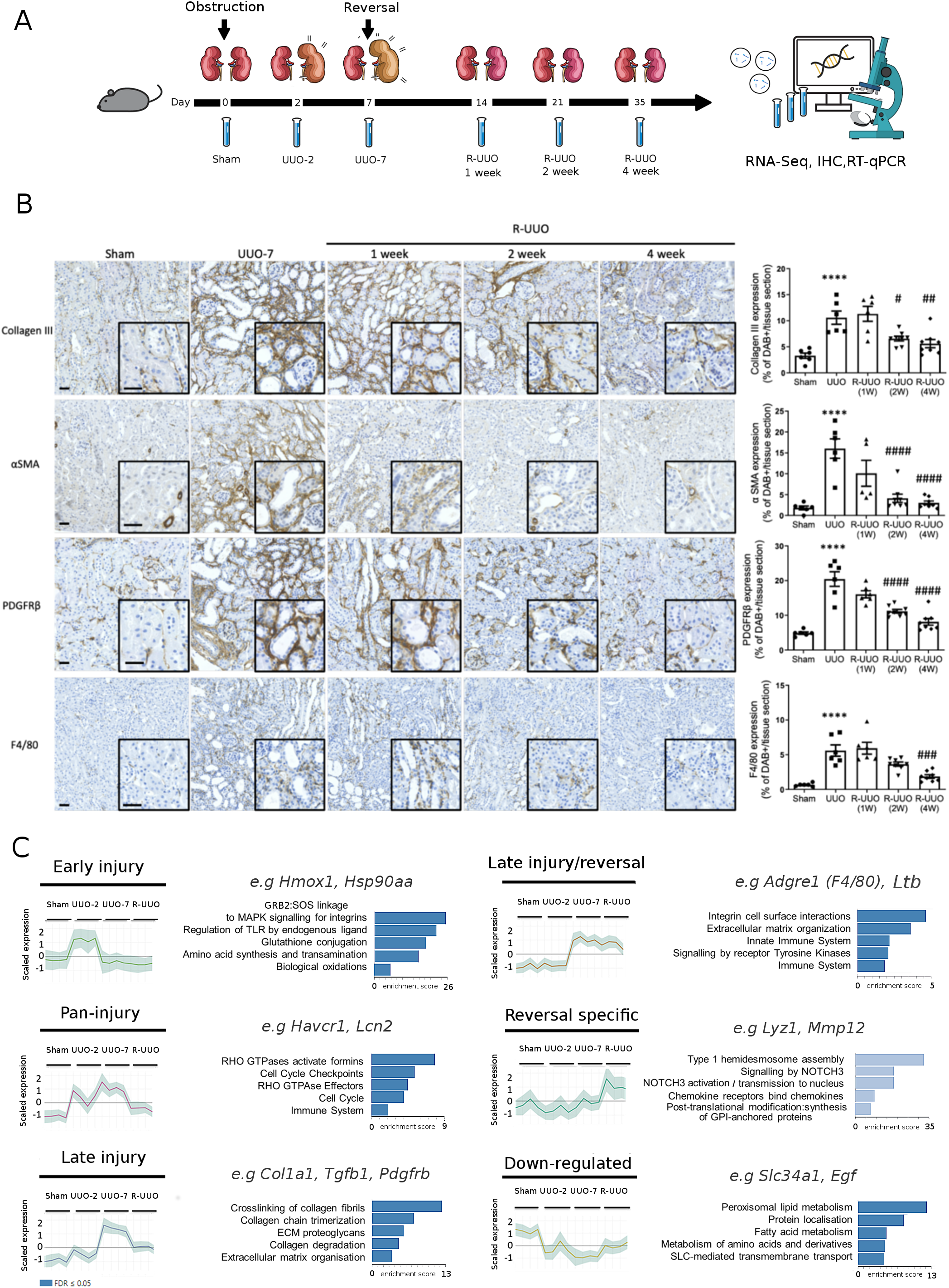
Characterisation of the reversible UUO (R-UUO) model. **a** Male 6-8 week old C57BL/6 mice underwent either UUO or sham surgery and were either culled 2 days later or left obstructed for 7 days and then culled or had their ureter re-implanted to reverse obstruction before sacrifice at 1, 2 or 4 weeks post R-UUO (n=6-8/group) **b** Representative images and quantification of fibrosis (collagen III) and fibroblast (PDGFR-β), myofibroblast (α-SMA) or macrophage (F4/80) accumulation during the R-UUO model. Scale bar 50μM ****P<0.0001 vs Sham, #p<0.05 vs UUO, ##p<0.01 vs UUO, ###p<0.001, ####p<0.0001 vs UUO. **c** Unbiased clustering analysis of bulk RNA sequencing data from the renal cortex of mice (n=4/group) during the R-UUO time-course identified 6 discrete temporal patterns of gene expression. Representative genes and enriched pathways are provided for each cluster. The number of genes included in each cluster is: 1: 1562, 2:779, 3:1479, 4:1619, 5:1708; 6:663. Shaded error range is the standard deviation of the mean scaled gene expression for each animal. Dark and light blue pathways are those demonstrating gene enrichment at a false discovery rate of <0.05 and >0.05, respectively.

### Kidney digestion for flow cytometry and single cell sequencing

Immediately after culling, mice were perfused with 10 mL PBS. Kidneys were excised, decapsulated and placed in ice-cold PBS. Equal portions of renal cortex from each mouse were finely minced in digest buffer [4.25mg/mL Collagenase V (Sigma-Aldrich St. Louis, Missouri, USA), 6.25mg/mL Collagenase D (Roche, Basel, Switzerland), 10mg/mL Dispase (Thermo-Scientific, Waltham, Massachusetts, United States) and 300μg/mL DNase (Roche, Basel, Switzerland) in RPMI 1640 (10% fetal calf serum, 1% penicillin/streptomycin/L-glutamine)] prior to homogenisation in gentleMACS^™^ C-tubes using the gentleMACS^™^ dissociator (Miltenyi Biotec, Auburn, California, USA). Samples were incubated at 37°C with shaking to maximise digestion. The kidney suspension was then subjected to a second gentleMACS TM homogenisation and digestion neutralised with an equal volume of FACS buffer (PBS, 2mM EDTA and 2% FCS). Kidney cell suspensions were then passed sequentially through 100 μm, 70 μm and 40 μm sieves. Any residual red blood cells were lysed by RBC lysis buffer (Sigma-Aldrich St. Louis, Missouri, USA). Cells were re-suspended in ice-cold FACS buffer ready for use.

### Flow Cytometry

Single-cell suspensions were incubated with FC block (BD Biosciences, San Jose, California) and then incubated with pre-conjugated antibodies (Suppl. Table 3) in round-bottomed plates. Controls were set up including unstained cells, beads with single stains of each antibody and FMOs (fluorophore minus one). DAPI was used to distinguish live and dead cells. For cellular composition analysis the following antibodies were used (CD45, CD31, LTL, PDGFRß, F4/80). For SMART-seq2 experiments, cells were incubated with the following antibodies (CD45, CD11b, CD11c, F4/80, MHCII, CD206, CD64, CD24, with a dump gate including TCRβ (T-cells), CD19 (B-cells), Siglec-F (eosinophils), Ly6G (neutrophils) all conjugated to BV421) and run on the BD FACS ARIA II. For paired blood exchange, blood and tissue were analysed on the BD 6L LSRFortessa using the following antibody panels: (Blood – CD45.1, CD45.2, CD3, F4/80, GR-1, CD11b, CD19; Kidney tissue – CD45.1, CD45.2, F4/80, Ly6C, MHCII, CD11b, CD206, CD24, CD64, and a dump gate including TCRβ, CD19, Siglec-F, Ly6G). All files were exported in FCS format and analysed with FlowJo software v10.

### Single-Cell droplet library preparation

For scRNAseq analysis on the 10x Genomic platform, single-cell suspensions from renal cortex were prepared from pools of 3 animals from each group as outlined by the 10x Genomics Single Cell 3’ v2 Reagent kit user guide. 50,000 live (DAPI-) cells were sorted on the BD FACS ARIA II. Samples were washed twice in PBS (Sigma) followed by centrifugation at 500g for 5 minutes at 4°C. Sample viability was assessed using trypan blue (Sigma) with an automated cell counter (Bio-Rad) and the appropriate volume for each sample was calculated. The chip was loaded with 10700 cells per lane.

After droplet generation, samples were transferred onto a pre-chilled 96-well plate, heat-sealed and reverse transcription was performed using a c1000 touch thermal cycler (Bio-Rad). After reverse transcription, cDNA was recovered using the 10x Genomics Recovery Agent and a Silane DynaBead cleanup (ThermoFisher) was performed. Purified cDNA was amplified and cleaned using SPRIselect beads (Beckman). Samples were diluted at 4:1 (elution buffer (Qiagen)/cDNA) and an initial concentration check performed on a Qubit fluorometer (Invitrogen) to ensure adequate cDNA concentration. Final cDNA concentration was checked on a bioanalyzer (Invitrogen).

### SMART-Seq2 library preparation

For the SMART-Seq 2 experiment, we performed flow cytometry for n=3 animals/group and sequenced 1 animal/group. The flow cytometry patterns within each group were broadly similar, mitigating against the selected animal being unrepresentative of the group. A single live (DAPI-) EGFP+ (Csf1r+) cell was sorted into each well of 96-well plates and all fluorochrome information was recorded using the Index Sort capability of the BD FACS ARIA II. Equal numbers of cells from each time-point were sorted into each plate to reduce batch effect, with 192 cells/time-point included in total. Single cells were processed as previously described (38). Briefly, cells were lysed immediately in lysis buffer containing 5% RNase inhibitor and 0.025% Triton X. Oligo (dT) primers were added and reverse transcription was performed. SMART-seq2 libraries were prepared according to the previously described protocol (38) with a few modifications (39): at step 5, 0.1 μl of External RNA Controls Consortium (ERCC) spike-in mix (10-5 diluted; Life Technologies, 4456740) was added with 0.1 μl of 100 μM oligo-dT primer and 1 μl of dNTP mix and 0.8 μl of H2O, yielding the same concentrations of primer and oligo as originally reported. Fluidigm protocol (PN 100-7168 M1) was used for tagmentation library generation. Final cDNA concentration was checked on a bioanalyzer (Agilent).

### Sequencing

The 10x libraries were pooled and normalised by molarity before being sequenced across 4 lanes on a single Illumina flow cell. Sequencing was performed on an Illumina HiSeq platform with a target of ~350M PE reads/lane, giving ~525M PE reads/sample comprising of 2×150bp PE configuration and 8bp index reads. The SMART-seq2 libraries were sequenced as 8x 96-plates which were pooled and sequenced on an Illumina HiSeq 4000 (50 bp, single-end reads). Data was deposited in the National Center for Biotechnology Information Gene Expression Omnibus database (accession #GSE140023)

### Single-cell RNA sequencing analysis

For the droplet-based dataset, the cellranger mkfastq wrapper (Cell Ranger Single Cell software suite 2.1.0, http://10xgenomics.com) de-multiplexed the illumina output BCL files to library specific FASTQ files. Subsequently, alignment was performed using the cellranger count function using STAR aligned 2.5.1b (40) against the Ensembl mouse reference genome version GRCm38.68. Correction and filtering of cell barcode and unique molecular identifiers followed, and the retained barcodes were quantified and used to generate a gene expression matrix. Summary sequencing statistics are provided in Suppl. table 4. The SMART-seq2 raw reads were similarly mapped against the Ensembl mouse reference genome version GRCm38.68 using STAR RNA-seq aligner (40) with the additional inclusion of the sequences for the ERCC spike-ins.

For our droplet-based dataset, a standard sequence of filtering, highly variable gene selection, dimensionality reduction and clustering were performed using the scRNA-seq analysis R package Seurat(v2.3.4)(41). Following alignment and initial pre-processing, we began our R workflow with 15,046 genes across 7,073 cells in our Sham group, 16,450 genes across 5,088 cells in our UUO-2 group, 17,368 genes across 7124 cells in our UUO-7 group and 17,227 genes across 6,096 cells in our R-UUO group. To exclude low-quality cells in both single-cell experiments, we then filtered cells that expressed fewer than 300 genes per 500 unique molecular identifiers, and to exclude probable doublets, cells with >10,000 unique molecular identifiers and <3,000 genes were removed. We used a mitochondrial filter to remove cells in which >50% of genes were mitochondrial, consistent with other renal-specific scRNA-seq projects (42, 43). This is a higher filter than has been used in non-renal single-cell analysis but reflects the high mitochondrial content in renal tubular epithelial cells. Any gene not expressed in at least 3 cells was removed.

For the SMART-seq2 data, identical metrics were used as above, but the mitochondrial filter threshold was lowered to 25%, and cells with >25% of reads mapping to the ERCCs were additionally excluded. Following filtering and QC, 15,046 genes across 4,540 samples of Sham mice, 16,450 across 3,101 samples of the UUO-2 mice, 17,368 genes across 5,563 samples of UUO-7 mice, 17,227 genes across 4,308 samples of the R-UUO mice and 13,517 genes across 362 samples in the SMART-seq2 data were taken forward for analysis resulting in 92 cells in the sham group, 102 in UUO-2, 103 in UUO-7 and 65 cells from the R-UUO group.

Normalisation was performed using the Seurat package to reduce biases introduced by technical variation, sequencing depth and capture efficiency. We employed the default global-scaling normalisation method “logNormalize” which normalised gene expression per cell by the total expression and multiplies the result by a scaling factor before log transformation. We then scaled the data and regressed out variation between cells due to the number of unique molecular identifiers and the percentage of mitochondrial genes.

The expression matrix subsequently underwent dimensionality reduction using principal component analysis (PCA) of the highly variable genes within the dataset. Using Seurat’s FindVariableGenes function (and computed using the LogVMR argument) we used log-mean expression values between 0.0125 and 3, and a dispersion cut-off of 0.5 to select genes. PCA was performed using these selected genes and 20 principal components were identified for subsequent analysis in each dataset, selected both visually using the elbow point on the bowplot and via the Jackstraw method.

Further cluster-based quality control was performed in the droplet data using the density-based spatial clustering algorithm DBscan which was used to identify cells on a tSNE map. We initially set an eps value of 0.5 and removed clusters with fewer than 10 cells. The remaining cells were then clustered again with eps value 1, followed by removing the clusters with fewer than 20 cells. Of note this allowed identification of a cluster characterized by high expression of heat shock genes including *Fos, Jun, Atf3*. This cluster was removed as it was considered to be an artefact of cell stress due to the experimental protocol as recently described (44). This procedure removed 158 (3.4%) from a total of 4,540 cells in sham mice, 137 (4.4%) of 3,101 cells in UUO-2 mice, 167 (3%) of 5,563 cells in UUO-7 mice and 83 (19%) of 4,308 cells in R-UUO mice.

Clusters were then identified using Seurat’s FindClusters function, built using the first 10 principal components and a resolution parameter of 1.5. The original Louvain modularity optimization algorithm was employed. tSNE (using the Rtsne package Barnes-Hut implementation) was then used for further dimensionality reduction and visualisation, which was run on a reduced dimensional space of the first 5-10 dimensions, using perplexity values of 15-50.

For all single-cell differential expression tests, we used the Wilcoxon rank-sum test to identify a unique expression profile for each cluster, with differential expression tested between each cluster and all other clusters combined. The FindAllMarkers test as implemented in Seurat returns an “adj_pval” (Bonferroni adjusted p-values) and an “avg_logFC” (average log fold change) for each gene. Genes were ranked in order of average log fold change and visualised using heatmaps.

The process of acquiring the myeloid subsets in Figure 3 required a combination of cluster-based cell pruning and a gene-based cell filter. The clusters were annotated by cell type, using the differentially expressed marker genes generated, and recognised markers based on known biology and on the single cell literature to date. All myeloid clusters were then isolated, renormalized and rescaled as described above, and were processed again through the pipeline described above, although without the DBscan QC step. Clustering resolution was lowered for each downstream implementation of FindClusters function and again clusters were identified based on differentially expressed genes. Any non-myeloid cell clusters were removed, and the data was once again, reprocessed and re-clustered. This entire process was repeated three times in total, with successively higher resolutions used to generate greater numbers of clusters, to coerce any non-myeloid cells into forming distinct clusters which could be removed as the data became cleaner. The expression of 52 key non-myeloid genes was then assessed and any cell expressing such genes was removed. Finally, tSNE graphs were visually inspected, and any unusual clusters were manually selected and DE genes inspected - any stray unwanted cluster was manually pruned and the data reprocessed as before.

**Fig. 2.**
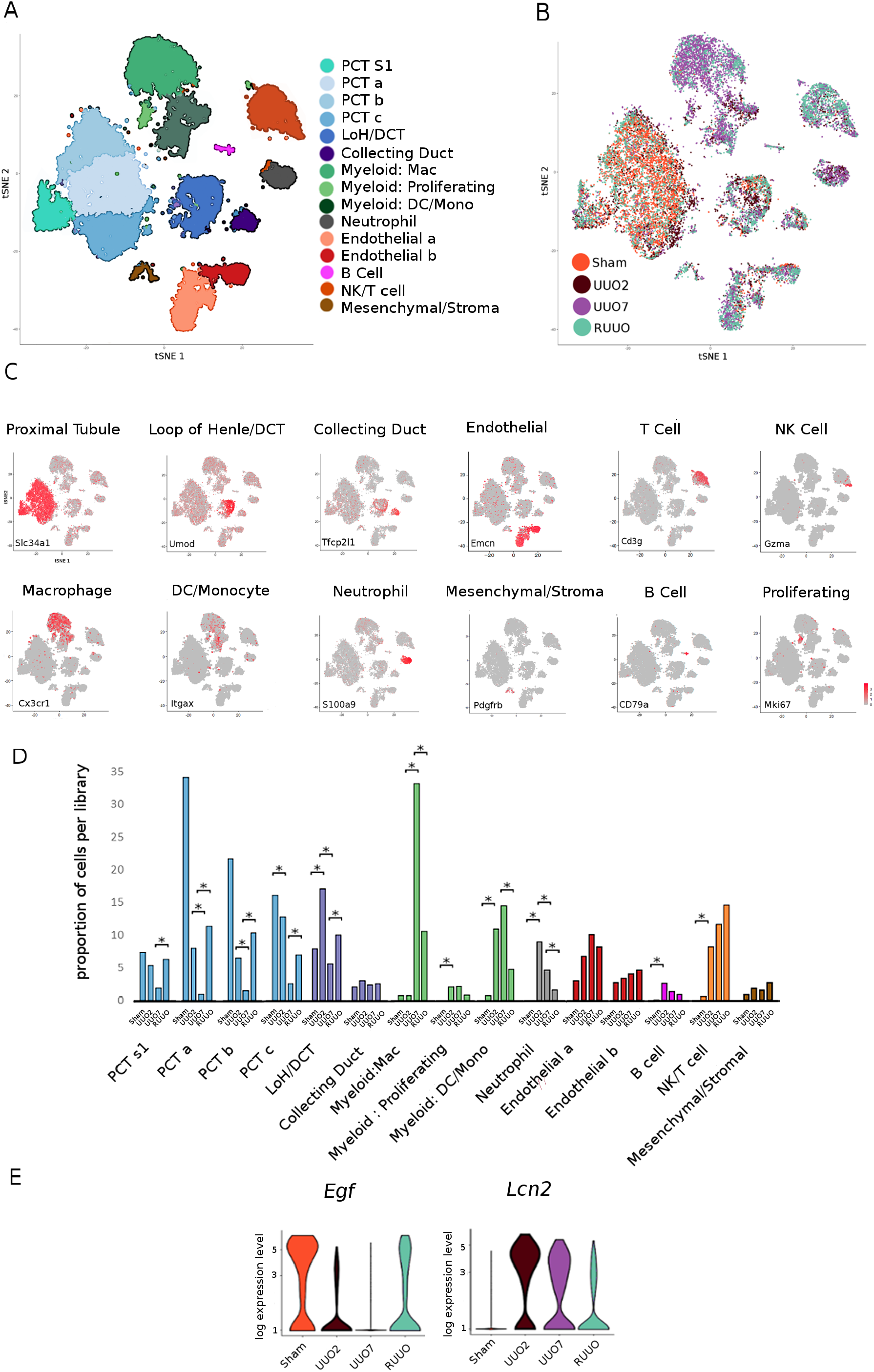
Single cell RNA-seq analysis of the kidney during R-UUO. tSNE plots of 17136 cells from libraries pooled from mice that underwent sham, UUO-2, UUO-7 or R-UUO (n=3/time-point) classified by **a** cell cluster and **b** time-point. **c** Expression of selected marker genes for each cell classification projected onto tSNE plot. Colour scale is Log10 expression levels of genes. **d** Relative proportions of cells assigned to each cluster by timepoint. Statistical significance derived using differential proportional analysis with a mean error of 0.1 over 100,000 iterations, *p-value<0.05. **e** Violin plots of *Egf* and *Lcn2* (encodes NGAL) gene expression in the Loop of Henle/Distal convoluted cell cluster. The y-axis shows the log-scale normalized read count. PCT: proximal convoluted tubule; S1: S1 segment; a-c: PCT sub-clusters coloured by shared nearest neighbour; LoH: loop of Henle; DCT: distal convoluted tubule, Mac: macrophage; DC: dendritic cell; Mono: monocyte, NK: Natural killer cell

**Fig. 3.**
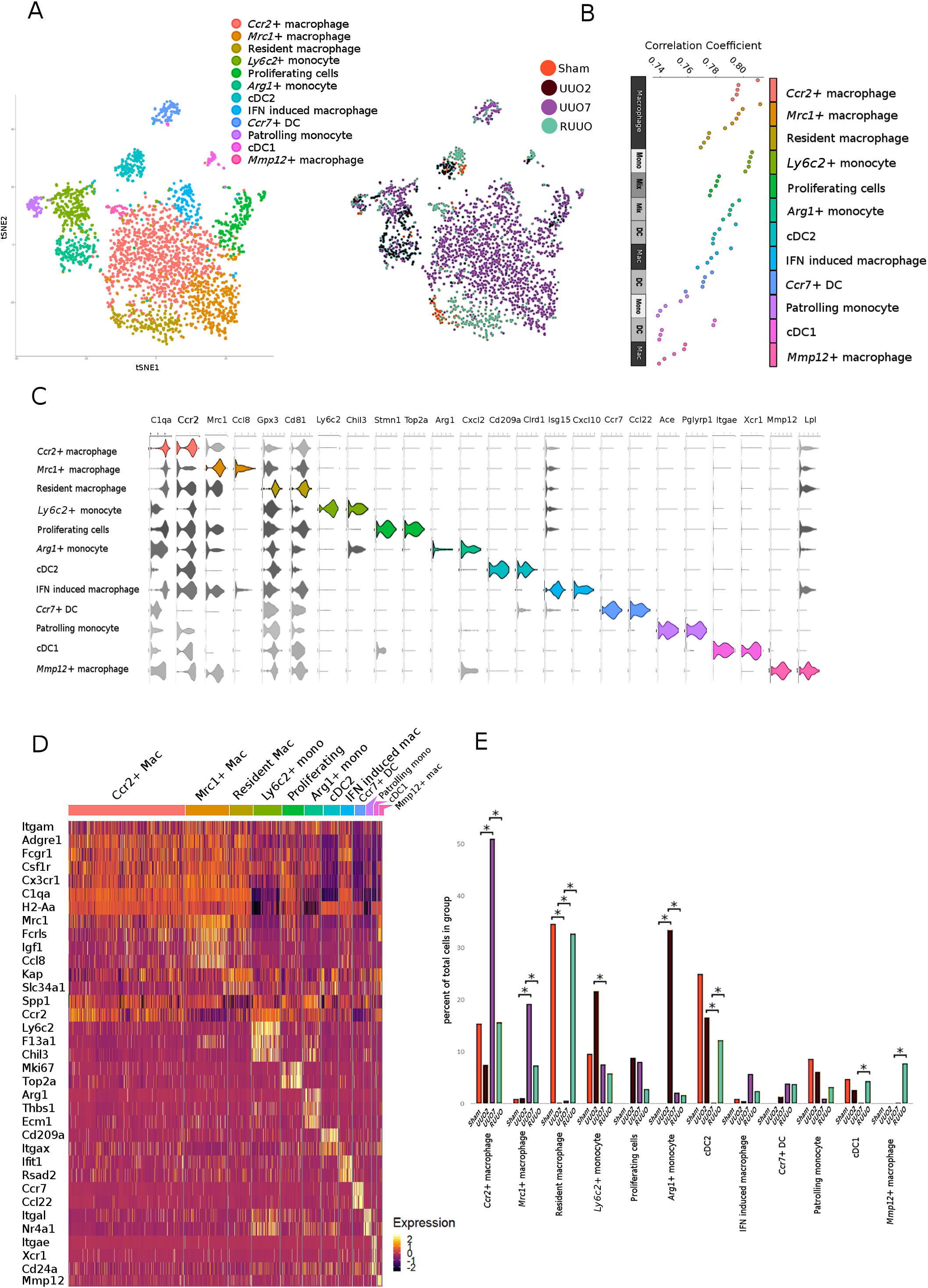
Single cell RNA-seq analysis of the myeloid cell populations. **a** tSNE plot of 2956 pooled myeloid cells from each time-point annotated by cell type and time-point. **b** Top 5 Immunological Genome Project (ImmGen) reference immune cell types correlating with each cluster, ranked by Spearman’s correlation coefficients following a cluster-to-references analysis using cluster identify predictor v2. **c** Violin plots showing the expression levels of selected marker genes in each cluster. The x-axis shows the log-scale normalized read count. **d** Heatmap of selected marker gene expression in each cluster calculated using Wilcoxon signed-rank test. The colour scheme is based on z-score distribution. **e** Relative proportions of each cell type at each time point. Statistical significance tested using differential proportional analysis with a mean error of 0.1 over 100,000 iterations, *p-value<0.05

### Differential Proportion Analysis

Differential proportion analysis (DPA) was developed to detect changes in population proportions across groups within single cell experiments. The algorithm is described in detail, along with source code by Farbehi *et al*. (45). Briefly, this approach employs a novel permutation-based statistical test to analyse whether observed changes in proportions of cell populations were greater than expected by chance. This approach attempts to consider sources of technical variation within the experimental technique (such as differing absolute cell numbers within the experiment, cell-type capture bias - a known feature of current single cell workflows (46) and variation due to *in silico* analysis (cluster assignment accuracy for example). A proportion table of clusters per phenotype/group is created from the count table, and the difference in cluster proportion is compared to a null distribution. This distribution is constructed by random permutations of random sub-samples of cluster labels across a random proportion of total cells. A new proportion table is generated from these data, and the process is repeated multiple times, with the resulting difference in cluster proportions across the data forming the null distribution. The observed distribution is then compared to this null-distribution and final p-values are calculated based on the minimum p-values of any observed increases or decreases in proportion. As per the original paper we used a w parameter= 0.1 where lower values will trend towards a stricter test (fewer significant hits) and higher values trend towards higher numbers of significant hits.

### Assignment to myeloid cell phenotype on ImmGen Consortium

Cluster identity predictor (v2) (https://aekiz.shinyapps.io/CIPR_v2/) was used to generated Spearman’s correlation values for each cluster (Fig. 3b) within our own data as compared to cluster gene signatures with Immunological Genome Project (ImmGen) mouse immune cell datasets based on the entire gene expression dataset. The algorithm first subsets genes common to both datasets before employing a one-to-many (cluster-to-references) calculation of correlation coefficients. A single correlation coefficient is calculated for each reference cell type and for each cluster allowing each cluster in our experiment to be analysed against each known cell type in the reference file and scored for its overall similarity. To further validate these assignments, we used SingleR (26) to assign myeloid cell classification using the murine Immgen dataset and create a consensus matrix with our classifications. Briefly, this pipeline is based on correlating reference bulk transcriptomic data sets of pure cell types with single-cell gene expression, similar to the Cluster identity predictor - a Spearman coefficient was calculated for single-cell gene expression with each of the samples in the reference data set based only on the variable genes in the reference data set. This is performed iteratively until a classification is reached. (Suppl. Fig 3c).

### Ligand-pair interactions

Heatmaps and dotplots of number of ligand-pair interactions were generated using the CellphoneDB tool (https://www.cellphonedb.org/) developed by the Teichmann Lab, Wellcome Sanger Institute, Cambridge, UK (47). The lower cut-off for expression proportion of any ligand or receptor in a given cell type was set to 10%, and the number of permutations was set to 1000. The clusters were not subsampled.

### Platform integration

As our SMART-seq2 library included an index sort, the transformed FACS data corresponding to each cell was then exported as .fcs files and analysed using FlowJo software. The index sort data was then extracted using the “index” FlowJo plugin available from FlowJo exchange. These data were then matched to the cell barcode and imported as both a separate “protein” assay and as metadata into the Seurat object to allow for visualization.

Our droplet-based and SMART-seq2 datasets were then integrated using the “anchoring” approach introduced in Seurat v3 (48). Here we create an integrated reference dataset and transfer the cell type labels onto our SMART-seq2 data. Briefly, this approach requires identification of “anchors” between the datasets which represent shared biological states. This involves jointly reducing the dimensionality of both datasets using diagonalized canonical correlational analysis before searching for mutual nearest neighbours in the new shared space. The paired cells are treated as anchors which represented shared biology across the datasets. Anchors were identified using the default parameters of the “FindIntegrationAnchors” function, with the argument dims=1:20.

### Pseudotime analysis

Lineage reconstruction and pseudotime inference was performed using the slingshot package (49). This method works by learning cluster relationships in an unsupervised manner and constructing smooth curves representing the lineage across 2 visualised pseudotime dimensions. Briefly, this involved first creating the raw expression matrix of the subsetted cells which were classified as “*Arg1*+ monocyte”, “*Ly6c2*+ monocyte”, “*Ccr2*+ macrophage by Seurat. This was followed by filtering genes not expressed in any cluster with <10 cells and having at least 3 reads within that cluster. Full quantile normalisation was then performed before dimensionality reduction using diffusion maps via the destiny package (50). Next cells were clustered to allow slingshot to infer a global structure of the lineage using a Gaussian mixture modelling implemented in Mclust (51) revealing 8 underling clusters in the data. Slingshot then constructed the cluster based minimum spanning tree and fitted the principal curve.

### Pathway Analysis

Gene set enrichment analysis (GSEA) and over representation analysis (ORA) were used to identify enriched pathways based on the differential expressed genes using WebGestalt (http://www.webgestalt.org/). For GSEA, we generated a rank for each gene in the list of DE genes using the formula (average: (average log fold change)*(-log (adjusted p-value)). To perform ORA of bulk RNA-Seq data, significantly DE genes were selected for the algorithm based on a minimum of 2-fold up-regulation (or 50% of baseline for the down-regulated genes) against the appropriate comparator (e.g. UUO2 v sham when considering early injury).

Enrichment categories were discarded if they contained less than 5 or over 2000 genes. These thresholds were calculated by WebGestalt based on the number of overlapping genes between the annotated genes in the category and the reference gene list for the ORA method. For the GSEA method, categories were discarded if they contained less than 15 genes or more than 500 genes. The Benjamini-Hochberg method was used to correct for multiple testing during ORA, and the top ten enriched categories as ranked by false discovery rate were selected. The reference gene list used was the Illumina mouseref 8. We used pathway gene sets from the protein analysis through evolutionary relationships, PANTHER (http://www.pantherdb.org), Reactome and Kyoto encyclopaedia of genes and genomes (https://www.genome.jp/kegg/) as our reference gene lists.

### Statistical analysis

Animal group size was determined from previous pilot experiments. Comparisons between two unpaired, non-normally distributed data points were carried out via Mann-Whitney test. Comparisons between two unpaired, normally distributed data points were carried out via T-test. Comparisons between multiple groups were performed with one-way ANOVA with Tukey’s multiple comparison test. All statistical analysis was performed using GraphPad Prism, version 10.

## Results

### Degradation of excess ECM following reversal of ureteric obstruction is associated with persistence of immune cells

To determine the pathways that mediate renal injury and repair, we utilised the murine R-UUO model (Fig. 1a). We performed immunohistochemistry and RNA sequencing (RNA-seq) on bulk renal cortical tissue from animals that underwent sham surgery, animals that had UUO surgery for 2 days or 7 days previously (hereafter referred to as UUO-2 and UUO-7, respectively) or those that had their ureter re-implanted after 7 days of obstruction (R-UUO) and were culled 1, 2 or 4 weeks later.

Within 7 days of ureteric obstruction there was expansion of interstitial PDGFR-β^+^ cells, activation of a proportion of these cells to myofibroblasts as determined by co-localisation of α-SMA^+^ and PDGFR-β^+^ (Suppl. Fig 1a) and collagen deposition (Fig. 1b), in association with marked induction of tubular injury markers such as *Havcr1* (encodes kidney injury molecule-1, KIM-1, Fig. 1c). Following re-implantation of the ureter, there was a decline in *Havcr1* expression and a significant reduction in interstitial PDGFR-β^+^ cells with reversal of myofibroblastic activation as evidenced by loss of PDGFR-β^+^α-SMA^+^ co-localisation (Suppl. Fig. 1a). Myofibroblast deactivation has not previously been described following R-UUO, but a similar phenomenon has been reported in the liver following cessation of injury (52). There was a more gradual regression of collagen deposition over 4 weeks as has been observed previously (35). There was evidence of innate immune system activation, with F4/80^+^ macrophages accumulating in the kidney during UUO, and persisting through the early stages of R-UUO, before trending towards control levels after 4 weeks of reversal (Fig. 1b) consistent with previous findings in this model which showed elevated numbers of macrophages were observed in the focal areas of the kidney being remodeled after R-UUO (35).

Analysis of the renal transcriptome has not previously been performed in the R-UUO model, therefore we performed unsupervised clustering of the RNA-seq data, which revealed 6 discrete temporal patterns of gene expression (Fig. 1c, Suppl. Fig. 1b-d, Suppl. Table 5). 3 of the clusters were characterized by gene up-regulation predominantly during the injury (UUO) phase with reversion towards baseline after R-UUO: i. genes up-regulated in early injury (UUO-2 only) were enriched for damage-associated molecular pattern (DAMP)-TLR signalling, MAPK signaling and oxidative stress pathways (Fig. 1c); ii. genes induced pan-injury included cell cycle genes and markers of kidney injury including *Havcr1* and *Lcn2* (encodes neutrophil gelatinase-associated lipocalin, N-GAL); iii. genes induced in late injury (predominantly restricted to UUO-7) were enriched for extracellular matrix (ECM) components, ECM cross-linkers and inhibitors of ECM degradation.

We identified 2 clusters which were characterised by gene activation predominantly during R-UUO and were enriched for genes implicated in innate and adaptive immunity (Fig. 1c). Remarkably, 5 of the top 10 genes induced specifically following R-UUO (*Lyz1, Mmp12, Gpnmb, Ccl8* and *Retnla*, Suppl. Fig. 1d,e) were also induced in macrophages in our model of resolution of liver fibrosis (31). In a final cluster, genes were down-regulated during UUO, partly reverted to baseline levels following R-UUO, and included markers of tubular cells from all segments of the nephron. Pathway analysis demonstrated enrichment for genes relating to tubular function including solute transport, protein/lipid metabolism, and detoxification (Fig. 1c).

The broad patterns of temporal gene expression in the R-UUO model were consistent with those we observed in a rat model of reversible diabetes and hypertension, with pro-fibrotic pathways induced specifically during injury, while immune pathways persist despite removal of the injurious stimuli (Suppl Fig. 1f,g) (32). Taken together, our current and previous data suggest common mechanisms of injury and repair across multiple models and organs, with regression of fibrosis being characterized by the presence of a specific macrophage phenotype.

### scRNAseq demonstrates dynamic changes in the proportion of intrinsic and immune cells during the R-UUO model

To characterise the heterogeneity of cells in the kidney during injury and repair and to assign the bulk transcriptomic changes to specific renal cell types, we performed scRNA-seq using the 10x Genomics platform on single-cell suspensions from the renal cortex of animals at four time-points: baseline (sham surgery), UUO-2, UUO-7 and 2 weeks after R-UUO (cells were pooled from 3 mice at each time-point). We used a shared nearest neighbour (SNN) approach to perform unsupervised clustering of the aggregated data from ~17,500 individual transcriptomes, identifying 15 discrete clusters (Fig. 2a, b, Suppl. Fig. 2a-d, Suppl. Table 6), which were assigned to renal cell types using known cell-specific markers in murine kidneys (Fig. 2c, Suppl. Fig 2b) (42). As this is the first scRNA-seq dataset of the kidney during and following reversal of chronic fibrotic injury we have generated an interactive online tool for the analysis of the dataset, available at https://argonaut.is.ed.ac.uk/shiny/katie.connor/mac_shiny/.

To analyse proportional changes of cell populations from injury to repair phases, we applied “differential proportional analysis”, a permutation based statistical method for detecting changes in population proportions across time points which considers sources of non-biological variation such as cell numbers, cell-type capture bias or cluster assignment accuracy (45). To complement this, we additionally used standard flow cytometry markers to analyse the proportional changes of key cell types within the kidney: proximal tubule cells (LTL), endothelial cells (CD31), fibroblasts/pericytes (PDGFR-b), immune cells (CD45) and F4/80hi and F480lo macrophages (Suppl. Fig.2e). Following UUO there was a marked reduction in the proportion of cells derived from the proximal tubule, which partly reversed following R-UUO (Fig. 2d, Suppl. Fig. 2e). While the proportion of cells derived from other tubular segments (Loop of Henle/distal convoluted tubule, collecting duct) was not reduced following UUO, there were dynamic changes in the tubular transcriptome, with induction of injury markers such as *Lcn2* (adj p-val 1.45E^-293^) and a reduction of biomarkers of tubular cell health such as *Egf* (adj p-val 2 5.67E^-05^) specifically during the injury phase (Fig. 2e) (53, 54). There was early recruitment of neutrophils and NK cells to the obstructed kidney, followed by expansion of the macrophage and T-cell populations, which persisted beyond the reversal of obstruction as observed previously during repair in a model of ischaemia-reperfusion injury (25) (Fig. 2d, Suppl. Fig 2e).

### scRNA-seq reveals myeloid heterogeneity during injury and repair

Our previous data in the kidney and liver suggest a pivotal role for the plasticity of myeloid cells in injury and repair (32, 47). To further characterise the myeloid cell heterogeneity and phenotype we repeated the SNN clustering specifically on myeloid cells, partitioning these cells into 12 clusters (Fig. 3a). To first assign unbiased broad classifications to these clusters, we generated Spearman’s correlation values for each cluster as compared to gene signatures of mouse immune cells obtained from the Immunological Genome Project (ImmGen) (Fig. 3b, Suppl. Fig. 3a). Similar results were obtained using the SingleR tool (Suppl Fig 3c). We then refined this classification manually using a combination of cluster-defining differentially expressed genes (Suppl. Fig. 3b, Suppl. Table 7) and genes that encode protein markers typically used to define specific myeloid cell subsets including *Itgam* (CD11b), *Adgre1* (F4/80), *Fcgr1* (CD64), *Itgax* (CD11c) and *H2-Aa* (MHCII) (Fig. 3c, d). Here, we were successful in describing key myeloid clusters during injury and for the first time during resolution of renal fibrosis at a single cell level.

### Early accumulation of *Ly6c2*^+^ and *Arg1*^+^ monocytes following ureteric obstruction

We uncovered two clusters that mapped to monocytes on ImmGen database (Fig. 3b), both of which expressed low levels of MHCII genes (*H2-Aa, H2-Ab1*), consistent with a monocytic phenotype (Fig. 3d). We identified the first cluster as patrolling monocytes as they expressed *Nr4a1* and *Itgal*, which are implicated in survival (55, 56) and adhesion (55, 57) of Cx3cr1^+^/Ly6C^-^ patrolling monocytes, respectively. As expected, the patrolling monocyte cluster did not expand in the kidney following injury (Fig 3a, e). We annotated the second cluster as inflammatory Ly6C^+^ monocytes as they expressed *Ly6c2, Ccr2, F13a1*, and *Chil3* genes (23). Cells in this cluster increased at UUO-2 (Fig 3a, e) and did not express the macrophage marker F4/80 as expected (Fig. 4a), indicating selective early recruitment of Ly6C^+^ inflammatory monocytes to the kidney during injury.

**Fig. 4.**
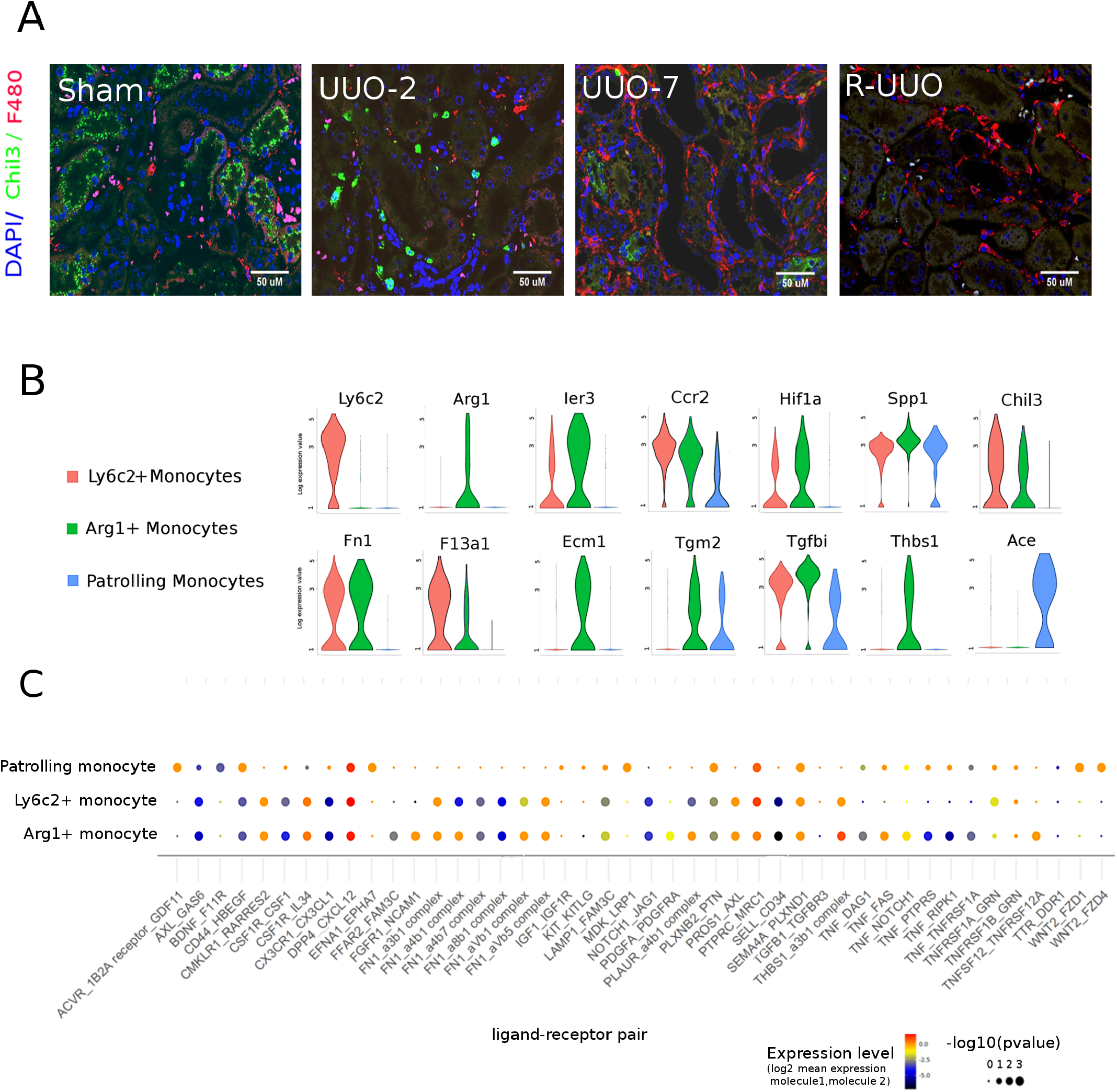
Characterisation of monocyte clusters. **a** Representative immunofluorescence images from each time-point in the R-UUO model for Chil3 (marker of Ly6C^+^ monocytes, red) and the pan-macrophage marker F4/80 (green). **b** Violin plots showing Log10 expression levels of selected genes in the 3 monocyte clusters. The y-axis shows the log-scale normalized read count. **c** Significant monocyte ligand - mesenchymal receptor pairs across the three monocyte subclusters. Colour of dot is proportional to mean expression values for all the interacting partners and size is inversely proportion to p-value.

Our single-cell clustering revealed a cell subset that mapped to a mixture of monocyte/macrophage states, but expressed low levels of MHCII, consistent with a monocytic phenotype (Fig. 3b). Cells in this cluster uniquely expressed *Arg1* (Fig. 3c, d) and although they did not express *Ly6c2*, they did express other markers of Ly6C^+^ inflammatory monocytes, including *Ccr2, Chil3* and *F13a1* (Fig. 3d, Fig. 4b). Differentially expressed genes in this cluster included early response genes (*Ier3, Fos, Jun*), hypoxia genes (*Hif1a, Vegfa*), pro-inflammatory genes (*Thbs1, Spp1*), pro-fibrotic genes (*Tgfb1, Tgfbi*), and genes encoding ECM components (*Fn1, Ecm1*) or ECM cross-linkers (*Tgm2*, Fig. 4b). The *Arg1*^+^ cells were exclusively present at UUO-2 (Fig. 3a, e) and their expression of pro-fibrotic genes suggested that they may initiate fibrosis by interacting with mesenchymal cells. Hence, we determined expression of ligand-receptor pairs between each monocyte subset and mesenchymal cells (Fig. 4c). More monocyte ligand-mesenchymal receptor pairs were expressed in *Ly6c2*^+^ and *Arg1*^+^ monocytes than patrolling monocytes. In addition, compared with the *Ly6c2*^+^ cells, the *Arg1*^+^ cells demonstrated greater potential for *Fn1-integrin, Pdgfa-Pdgfrβ* and *Tnf-Tnfsfr1* signalling to mesenchymal cells. Taken together, these data suggest that *Arg1*^+^ cells may be derived from recruited Ly6C^+^ monocytes that become activated acutely in the hypoxic and inflammatory milieu of the injured kidney towards a pro-fibrotic phenotype.

### Macrophages adopt differing phenotypes during injury and resolution phases

We identified 5 clusters mapping to macrophage phenotypes on the ImmGen database (Fig. 3b), all of which expressed genes consistent with macrophage identity including those encoding CSF1 receptor (*Csf1r*), MHCII (*H2-Aa*) and *Cd81*, a recently described marker of renal macrophages across multiple species (Fig. 3c, d) (23). We were able to functionally annotate many of these clusters.

We define one macrophage cluster as quiescent resident macrophages, as they were detected exclusively in kidneys from animals that underwent sham surgery or had their kidney de-obstructed (Fig 3a, e). They were tagged by markers of proximal tubular cells, which most likely represents the presence of ambient tubular RNA species, which is most prevalent in sham or R-UUO mice where the proportion of proximal tubular cells is greater than in UUO (Fig 2d). In contrast to other macrophage clusters, all of which expanded during UUO, these cells did not express *Spp1*, which encodes osteopontin, a marker of activated macrophages (58) that promotes renal injury following UUO (59).

We annotated second macrophage cluster as being involved in scar resolution and tissue regeneration. This cluster expressed high levels of *Mrc1* and cells from this cluster were most commonly observed at UUO-7 and to a lesser extent R-UUO (Fig 3a, e). *Mrc1* encodes mannose receptor (MR), and in keeping with this there was expansion of MR^+^ cells at in the renal interstitium at UUO-7 and co-localisation with the macrophage marker F4/80 (Fig. 5a). The *Mrc1*^+^ cells expressed multiple scavenger receptors (*Mrc1, Fcrls, Stab1*) suggesting a role in scavenging debris/excess ECM (Fig. 3d). In addition, they expressed *Igf1*, which is up-regulated in reparative macrophages in the liver (31) and promotes regression of cirrhosis and liver regeneration (60) and *Apoe* which dampens inflammation (61) and promotes regeneration (62). Hence the *Mrc1*^+^ macrophages expand in the later stages of injury, partially persist during resolution and adopt a transcriptomic profile consistent with degradation of scar and organ regeneration.

**Fig. 5.**
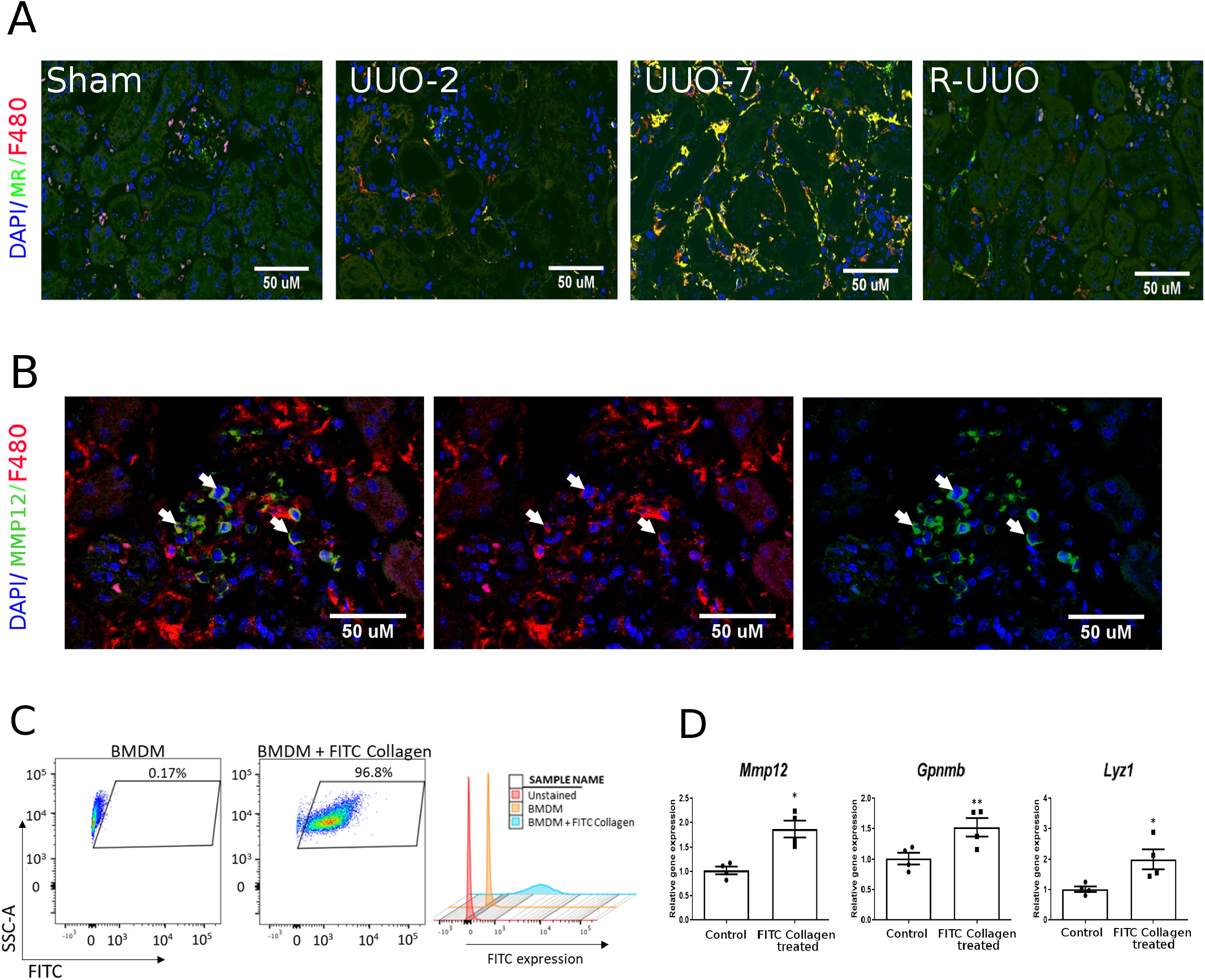
Characterisation of macrophage clusters. **a** Representative immunofluorescence images across the R-UUO time course for mannose receptor (MR, marker of *Mrc1*^+^ macrophages, green) and F4/80 (red). **b** Immunofluorescence for MMP12^+^ (green) and F4/80 (red) in a kidney 2 weeks post R-UUO. Arrows highlight MMP12^+^ cells that also stain weakly for F4/80 **c** Flow cytometry plots of bone marrow derived macrophages (BMDMs) demonstrating fluorescence following phagocytosis of FITC-collagen **d** Expression of reparative macrophage genes measured by qRT-PCR in BMDMs following phagocytosis of FITC-collagen v BMDMs cultured in medium alone (control). (n=4 replicates, *p<0.05 and **p<0.01).

A third macrophage cluster was predominantly comprised of cells from UUO-7 kidneys. In addition to macrophage markers, cells in this cluster expressed *Ccr2* (Fig. 3d), raising the possibility that they may be derived from Ly6C^+^/CCR2^+^ monocytes that are recruited to the injured kidney before transitioning to adopt a more macrophage-like phenotype.

A fourth cluster was uniquely characterised by expression of interferon-stimulated genes (Fig 3c, d). The function of these cells remains unknown, but a similar phenotype has been observed following injury in other organs, including the heart (27).

Intriguingly, a final macrophage cluster was observed solely in kidneys from mice that had undergone R-UUO and was characterised by expression of *Mmp12*, a macrophage-specific metalloproteinase, suggesting that these cells may be involved in matrix remodelling. Cells in this cluster expressed additional macrophage markers including *Cd81*, MHCII genes (*H2-Aa*), and scavenger receptors (*Mrc1, Fcrls*), but exhibited low expression of the *Adgre1* gene (Fig. 3d) and its protein product F4/80 (Fig. 5b). We have previously shown that *Mmp12*^+^ expression defines a reparative macrophage phenotype that mediates regression of liver fibrosis (46), and these *Mmp12*^+^ cells may have a similar role in kidney repair. Indeed, following phagocytosis of FITC collagen (Fig. 5c), bone marrow-derived macrophages up-regulated the degradative enzymes expressed in reparative macrophages including *Mmp12, Gpnmb* and *Lyz1* (31) (Fig. 5d) suggesting that they may switch to a matrix-degrading phenotype on encountering scarred matrix.

An additional myeloid cluster mapped to a mixture of cells in the ImmGen database (Fig. 3b) and expressed cell cycle genes, such as *Mki67* and *Top2a* (Fig. 3d), consistent with proliferation. Analysis of cluster-defining marker genes in the proliferating cells suggested that they were comprised largely of macrophages, with representation from all macrophage clusters (Fig. 3d).

In summary, we identified transcriptionally distinct macrophage phenotypes populating the kidney at specific times in injury and repair, highlighting the plasticity of this cell type.

### Dendritic cells adopt a migratory phenotype during late stage injury and resolution

We mapped 3 clusters to dendritic cells (DC) on the ImmGen database, all of which expressed MHC genes (*H2-Aa*) but not macrophage markers (*Cd81, C1q*, Fig. 3d). One cluster expressed *Itgae* (encodes CD103), while the other expressed *Cd209a*, consistent with type 1 and type 2 conventional dendritic cells (cDC1, cDC2), respectively (23). Both of these clusters were present in sham animals but were proportionally reduced by UUO-7 before returning during R-UUO (Fig. 3a, e). In contrast, the third DC cluster, which expressed *Ccr7*, was not detected in sham animals, but appeared by UUO-7 and persisted through R-UUO (Fig. 3a, e). This cluster mapped specifically to lymph node dendritic cells in the ImmGen database (Suppl Fig. 3a). Taken together the data suggest that following kidney injury resident DCs up-regulate *Ccr7* and this may promote migration to draining lymph nodes by binding to CCR19/CCR21 (63).

### Conventional flow cytometry does not capture the full heterogeneity of myeloid cells

To assess how our scRNA-seq-derived clusters corresponded to typical myeloid cell phenotypes observed using a conventional antibody approach on flow cytometry, we performed further scRNA-seq sequencing using plate-based SMART-seq2 technology which enables linking of the transcriptome in each cell to abundance of cell surface markers (FACS intensity) using an index sorting approach (64).

We repeated the R-UUO model using Macgreen mice (express EGFP under the Csf1r promoter) and performed flow cytometry, gating on CD45^+^Macgreen^+^TCRβ^-^CD19^-^Ly6G^-^Siglec-F^-^ myeloid cells. There was expansion of the CD11b^+^F4/80^Lo^ population at UUO-2, consistent with early recruitment of monocytes to the kidney in response to injury (Fig. 6a). By UUO-7, the monocyte recruitment had diminished but there was marked expansion of the CD11b^+^F4/80^Hi^ macrophage population, which persisted through 2 weeks following R-UUO (Fig. 6a).

**Fig. 6.**
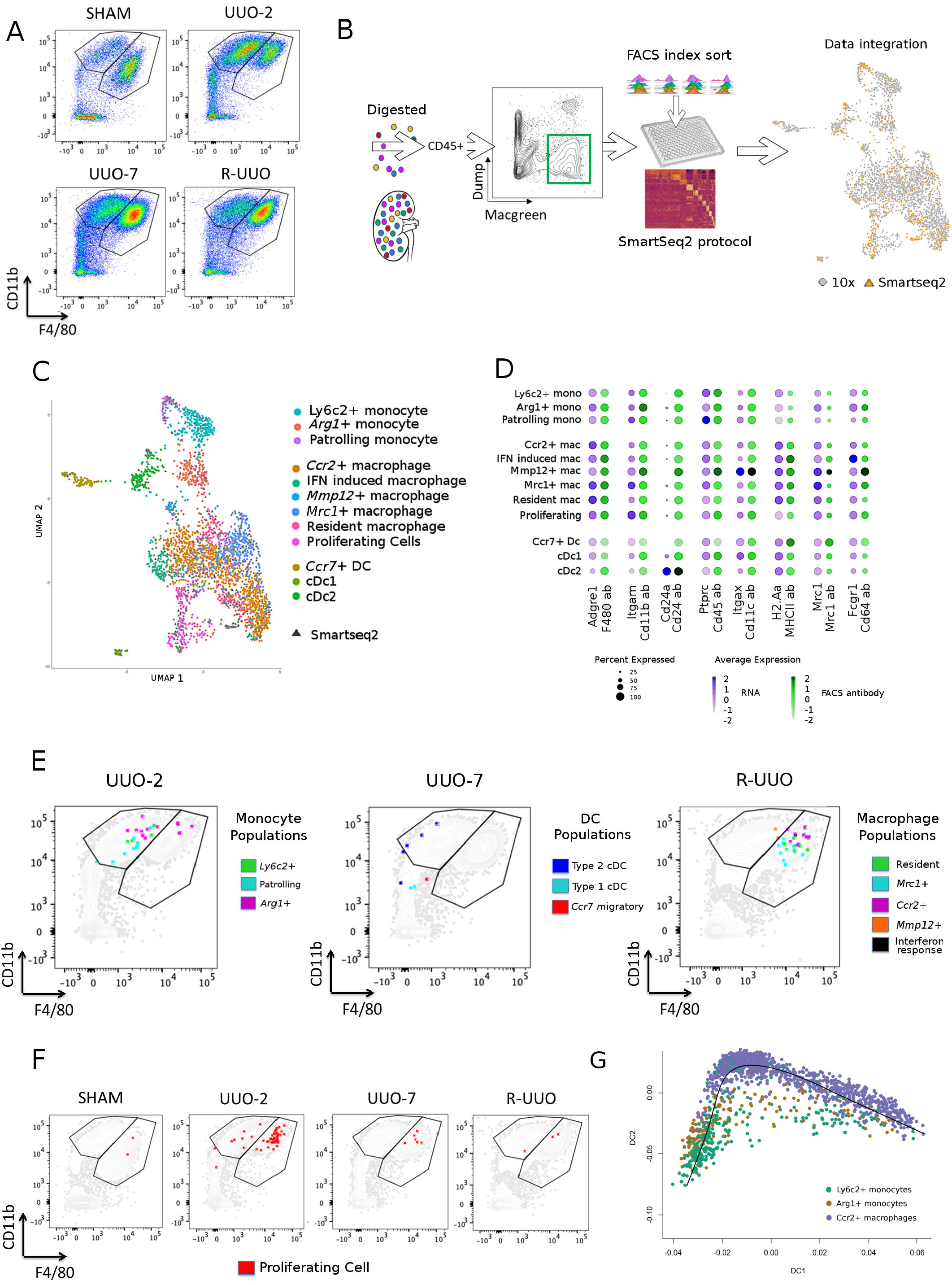
Mapping of the myeloid clusters to conventional flow cytometry using integrated droplet and plate based scRNA-seq datasets. **a** Representative flow cytometry plots from kidney cell suspensions from each time-point following gating on CD45^+^Macgreen^+^TCRβ^-^CD19^-^Ly6G^-^Siglec-F^-^ myeloid cells. Cells segregated into CD11b^+^F4/80^Lo^ monocyte and CD11b^+^F4/80^Hi^ macrophage gates **b** Strategy to integrate gene and cell surface protein expression at the single cell level. Kidneys were digested into single cell suspensions and single CD45^+^Macgreen^+^TCR1β^-^CD19^-^Ly6G^-^SiglecF^-^ myeloid cells were sorted into individual wells following index linkage to cell surface marker expression. They underwent scRNA-seq using the SMART-seq2 protocol before integration with the 10x dataset **c** UMAP of the combined 10x and SMART-seq2 dataset **d** Dot plot of cell surface protein and corresponding gene expression in each cluster. The size of the dot denotes the percentage of cells in each cluster expressing the relevant gene/protein; the intensity of colour represents mean gene/protein expression **e** Representative flow cytometry plots from UUO-2, UUO-7 and R-UUO illustrate mapping of cells from each myeloid cluster onto the CD11b^+^F4/80^Lo^ monocyte and CD11b^+^F4/80^Hi^ macrophage gates. **f** Mapping of proliferating cells (red) at each time-point onto the flow cytometry plots **g** Pseudotime analysis of the transcriptomes of the *Ly6c2^+^, Arg1*^+^ and *Ccr2*^+^ clusters

We sorted single CD45^+^Macgreen^+^TCR1β^-^CD19^-^Ly6G^-^Siglec^-^ myeloid cells into individual wells, capturing 192 cells from each time-point (sham, UUO-2, UUO-7 and 2 weeks following R-UUO, Fig. 6b). The cells underwent scRNAseq using the SMART-seq2 platform and this dataset was then integrated with the 10x dataset using a canonical correlation analysis derived “anchoring” method allowing the SMART-seq2 cells to be mapped to the myeloid cell identities derived from the 10x dataset. SMART-seq2 cells were observed in every cluster in the combined dataset (Fig. 6b,c). By employing indexing sorting alongside the SMART-seq2 platform we confirmed moderate correlation between the mean gene and corresponding surface protein expression observed within each cluster (Fig. 6d). We next mapped the cells from each myeloid cluster onto the monocyte and macrophage gates on flow cytometry (Fig. 6e). As not all clusters were represented at all time-points in the SMART-seq2 dataset, we used UUO-2, UUO-7 and R-UUO to illustrate mapping of cells from the monocyte, dendritic cell and macrophage clusters, respectively. Cells from the *Ly6c2*^+^ and patrolling monocyte clusters mapped to the CD11b^+^F4/80^Lo^ monocyte gate, as expected. Conversely, resident, *Mrc1*^+^, *Ccr2*^+^ and interferon-response macrophages all mapped appropriately to the CD11b^+^F4/80^Hi^ macrophage gate. Cells from the *Arg1*^+^ cluster straddled the monocyte and macrophage gates (Fig. 6e), with a proportion of the *Arg1*^+^ cells co-locating with *Ccr2*^+^ macrophages in the CD11b^Hi^F4/80^Hi^ region, suggesting that they may be transitioning to *Ccr2*^+^ macrophages. To assess this further, we performed pseudotime analysis, which revealed a transcriptomic trajectory consistent with transition of *Ly6c2*^+^ monocytes to *Ccr2*^+^ macrophages with *Arg1*^+^ monocytes representing an intermediate transitional state (Fig. 6f). Cells from the *Mmp12*^+^ cluster also straddled the monocyte and macrophage gates (Fig. 6e), consistent with their intermediate F4/80 expression on immunofluorescence (Fig. 5b). Cells from the cDC1 and *Ccr7*^+^ clusters were CD11b^-^F4/80^-^, whereas the cDC2 cells mapped to the CD11b^+^F4/80^Lo^ monocyte gate (Fig. 6e). Cells from the proliferating myeloid cluster were observed predominantly at UUO-2 and mapped to the monocyte and more particularly the macrophage gates (Fig. 6f). Macrophage proliferation was still present at UUO-7 and to a lesser extent R-UUO.

In summary, while cells defined as monocytes or macrophages using gene markers on scRNA-seq broadly localize to the appropriate monocyte/macrophage gate on flow cytometry, scRNA-seq detects additional myeloid heterogeneity, including the *Arg1*^+^, interferon-response and *Mmp12*^+^ clusters, which would not readily be detected by flow cytometry alone.

### Monocytes recruited early following UUO transition to a macrophage phenotype

In combination the pseudotime and flow cytometry data suggested that there was early recruitment of Ly6c^+^ monocytes to the obstructed kidney and that these may transition towards a *Ccr2*^+^ macrophage phenotype by UUO-7. To track monocyte fate following UUO, we performed paired blood exchange between Ly5.1 mice (CD45.1/CD45.2 heterozygous) and C57BL6/J mice (CD45.2 homozygous) (Fig. 7a). On day 1 following UUO, we performed 15 reciprocal exchanges of 150μl whole blood between each pair over a 20 minute period, resulting in ~40% of total CD45^+^ circulating cells being derived from the donor immediately after the exchange (Fig. 7b). The proportion of donor-derived circulating monocytes/neutrophils fell rapidly to ~1% by 2 days after UUO, with negligible numbers persisting in the circulation by 7 days (Fig 7c, full gating strategy in Suppl. Fig. 6a). There was a similarly rapid reduction in the proportion of donor-derived circulating T- and B-lymphocytes, however a small number of donor lymphocytes persisted in the circulation through UUO-7 (Fig. 7c).

**Fig. 7.**
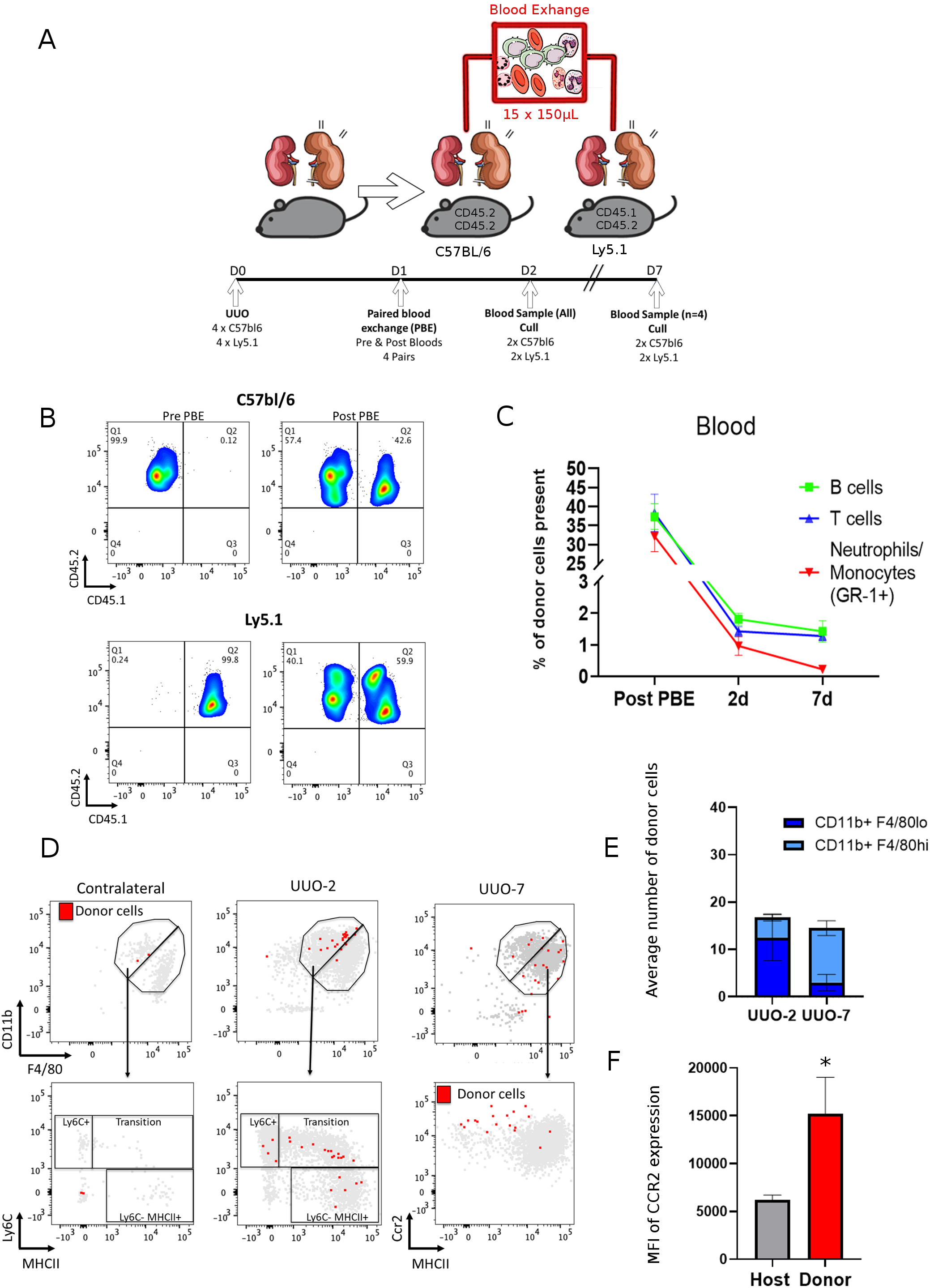
Tracking of monocyte recruitment to kidney post UUO by paired blood exchange. **a** Schemata of experimental strategy for paired blood exchange to track fate of immune cells recruited to the kidney. One day after UUO whole blood exchange was performed between pairs of C57BL/6 and Ly5.1 mice (n=4) and pairs were culled either 2 or 7 days after UUO **b** Representative flow cytometry plots of circulating CD45^+^ cells from pairs of mice pre- and immediately post-blood exchange illustrating approximately 40% of circulating cells were derived from donors after the exchange (CD45.1^+^CD45.2^+^ cells in C57BL/6; CD45.2^+/+^ cells in Ly5.1 mice) **c** Percentage of circulating immune cells derived from paired-donor over the experimental time-course (n=4 pairs immediate post PBE and at UUO-2; n=2 pairs at UUO-7) **d** Illustrative flow cytometry plots mapping donor cells (red) and recipient cells (grey) to the CD11b^+^F4/80^Lo^ monocyte and CD11b^+^F4/80^Hi^ macrophage gates, the monocyte ‘waterfall’ in obstructed and contralateral kidneys at 2 post UUO and the expression of CCR2 and MHCII 7 days post UUO. **e** Average number of donor cells mapping to the CD11b^+^F4/80^Lo^ monocyte and CD11b^+^F4/80^Hi^ macrophage gates in obstructed kidneys at 2 and 7 days after UUO. **F** The expression of CCR2 (MFI) on the donor cells compared to the host cells from the CD11b^+^F4/80^Hi^ macrophage gate in obstructed kidneys 7 days after UUO. * p<0.05 Mann-Whitney n=4/gp.

To determine the fate of donor monocytes recruited to the obstructed kidney, we performed flow cytometry on kidney cell suspensions, gating on CD45^+^CD64^+^TCRβ^-^CD19^-^Ly6G^-^SiglecF^-^ myeloid cells (Suppl Fig. 6b). At UUO-2, donor cells were recruited selectively to the obstructed kidney, with few observed within the contralateral kidney (Fig. 7d). As expected, the donor cells mapped almost exclusively to the CD11b^+^F4/80^Lo^ monocyte gate (Fig. 7e). Furthermore, the donor cells mapped to all areas of the monocyte ‘waterfall’, suggesting they were transitioning from a Ly6C^Hi^/MHC^Lo^ monocyte towards a Ly6C^Lo^MHCII^Hi^ macrophage-like phenotype in line with our previous observations (Fig. 7d). At UUO-7, some donor cells persisted in the CD11b^+^F4/80^Lo^ monocyte gate, but they were now predominantly Ly6C^Lo^MHCII^Hi^ suggesting a switch towards a more macrophage-like phenotype (Fig. 7d). Furthermore, the majority of donor cells at UUO-7 were located in the CD11b^+^F4/80^Hi^ macrophage gate, and they expressed high levels of CCR2 compared with the global macrophage population (Fig. 7f). Taken together, these data suggest that donor monocytes are recruited selectively to the obstructed kidney by UUO-2, transition to a CCR2^Hi^ macrophage by UUO-7, and hence are the likely source of the cells in the *Ccr2*^+^ macrophage cluster observed at UUO-7 in the scRNA-seq dataset (Fig. 3a).

### Myeloid cell subsets correlate with fibrosis in human kidney disease

To determine whether the myeloid cell phenotypes that we identified in murine obstructive nephropathy are also observed in human kidney disease, we assessed whether antibodies against myeloid cluster-specific markers bound to immune cells in the kidney using the Human Protein Atlas (Fig. 8a). Cells that stained with F13A1 (marker of *Ly6c2*^+^ monocytes) and DOK2 (*Arg1*^+^ monocytes) were located specifically in focal areas of injury/inflammation, whereas ITGAL (patrolling monocytes) was largely restricted to cells within the circulation. CD68, a pan-macrophage marker including resident macrophages, was widely distributed in the healthy kidney, whereas mannose receptor (*Mrc1*^+^ macrophages), and CCR2 (*Ccr2*^+^ macrophages) localized to areas of tissue injury. IRF8 and CD209, markers of types 1 and 2 conventional dendritic cells respectively, localised to areas of renal injury, with CCR7 (*Ccr7*^+^ migratory DCs) staining a cluster of cells that resembled a tertiary lymphoid follicle. To assess whether the cell-specific markers correlated with kidney disease, we employed RNA-seq data derived from renal biopsies taken from live kidney donors and from patients with diabetic nephropathy or focal segmental glomerulosclerosis recruited to the European Renal cDNA bank and freely available via the NephroSeq platform (www.nephroseq.org). The gene expression of each marker correlated with expression of *Col1a1* (encodes collagen I, Fig. 8b). Expression of *Mmp12* was not detected in the healthy human kidney or in patients with CKD, which is consistent with the fact that cells in the *Mmp12*^+^ cluster were specific to the resolution phase of kidney injury. The association between marker genes of murine myeloid subsets and *Col1a1* expression suggests that equivalent subsets may be present in humans, however more definitive evidence will be required from scRNAseq studies in human kidney disease.

**Fig. 8.**
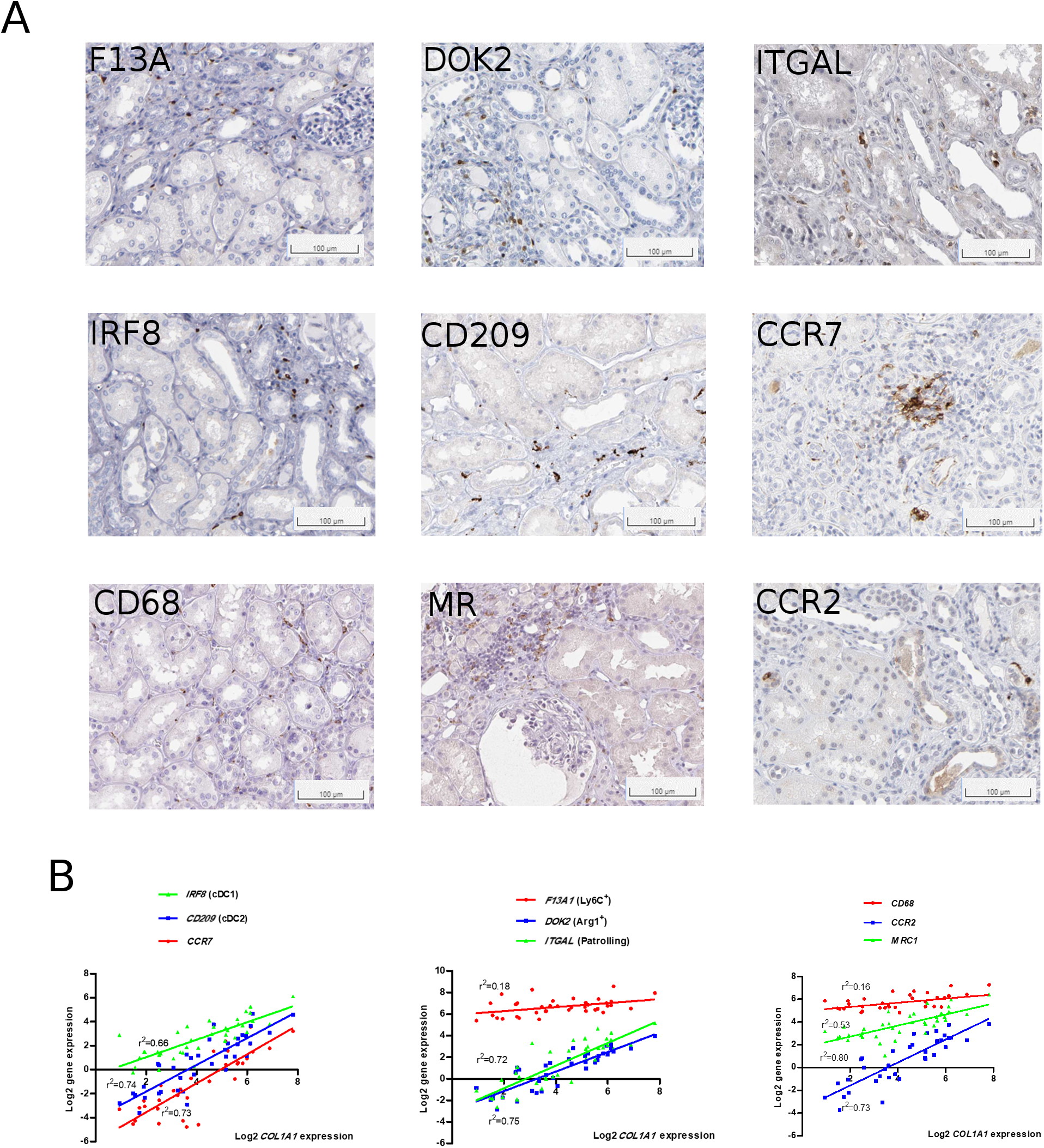
Markers of murine myeloid subsets in human kidney disease. **a** Immunostaining for myeloid cluster-defining markers in human kidney tissue obtained from the Human Protein Atlas (66). **b** Gene expression of cluster-defining markers for dendritic cells (left), monocytes (centre) and macrophages (right) against COL1A1 gene expression in the tubulointerstitium of kidneys from healthy controls (n=9), patients with diabetic nephropathy (n=10) and patients with focal segmental glomerulosclerosis (n=18). Data obtained from www.Nephroseq.org

## Discussion

Previous studies have assessed the kidney during repair following ischaemia-reperfusion injury, however our scRNA-seq studies represent the first detailed characterisation of the myeloid cell heterogeneity in the injured and repairing kidney, identifying novel monocyte and macrophage subsets not previously observed in the kidney. Acute injury induces a novel population of cells which are transcriptomically aligned to monocytes but which uniquely express *Arg1*^+^. While *Arg1* has traditionally been thought of as a marker of alternative macrophage activation (65), the *Arg1*^+^ cells do not express other markers of alternative activation such as *Mrc1* or MHCII-encoding genes, suggesting that *in vitro* immune activation assays do not reflect the complex *in vivo* milieu. Indeed, the *Arg1*^+^ cells express pro-inflammatory and pro-fibrotic genes, and future work should determine whether specific depletion of these cells could reduce disease severity. *Mrc1*^+^ macrophages expand in the late phase of injury and partly persist through the repair phase, with their transcriptome consistent with a role in scavenging debris and producing a pro-resolution secretome. Intriguingly, a novel *Mmp12*^+^ macrophage subset emerged specifically during the resolution phase. We have previously reported a similar macrophage phenotype during the resolution of liver disease, suggesting common reparative mechanisms across organs (31). Our *in vitro* studies suggest that ingestion of excess ECM or cell debris (31) may be a stimulus for induction of this phenotype, however their cellular origin and role in resolution requires further study as strategies to induce this phenotype may enhance scar degradation in the diseased kidney and other organs.

The cellular origin of the novel myeloid clusters detected during injury and repair is crucial for understanding their function and to develop therapeutic strategies. To track the fate of circulating immune cells recruited to the kidney, we employed paired blood exchange, which has previously been used to assess the effects of donor serum (37) but not, to our knowledge, to track immune cells. By combining paired blood exchange and flow cytometry, we demonstrate early recruitment of monocytes specifically to the obstructed kidney and that these subsequently adopt a macrophage phenotype, but continue to express CCR2. These results suggest that circulating monocytes are the source of the large *Ccr2*^+^ macrophage cluster observed at UUO-7 in the scRNA-seq dataset, which is consistent with lineage tracing and parabiosis studies following myocardial infarction (27). Remarkably, although they still express *Ccr2*, their transcriptome is otherwise almost identical to resident macrophages. In addition, genetic or pharmacological inactivation of CCR2 following renal ischaemia-reperfusion injury reduces the expansion of F4/80^+^ macrophages in the kidney and the severity of renal fibrosis, suggesting that CCR2^+^ cells may be detrimental (19). One advantage of paired blood exchange over parabiosis or bone marrow transfer is that the donor cells persist at large numbers in the circulation for a relatively short time, therefore enabling tracking of cells at multiple discrete time-points after injury or during resolution of disease. Future studies may provide insight into the source of the cells in the *Mmp12*^+^ cluster. The short circulating time of donor cells may also be a limitation of the technique, in that only a small proportion of recruited cells are derived from the donor, therefore a limited number of cells are available for downstream analysis. Further refinements including performing several consecutive PBE may increase the yield.

In summary, by combining complementary technologies, including plate and droplet-bsed scRNA-seq, flow cytometry and paired blood exchange, our studies have identified novel subsets of myeloid cells, which may offer therapeutic targets to inhibit progression and enhance resolution of kidney disease.

## Acknowledgements

We acknowledge Gary Borthwick, Centre for Cardiovascular Science, University of Edinburgh for his assistance in performing the R-UUO surgery. Flow cytometry data was generated with support from the QMRI Flow Cytometry and Cell Sorting Facility, University of Edinburgh. Thanks to Mike and Irina Conboy (University of California, Berkley) for assistance with establishing the paired blood exchange model.

## Authors’ contribution statement

Funding acquisition, supervision, conceptualization, project administration, methodology, investigation, analysis, writing and editing manuscript – LD, BRC. Funding acquisition, conceptualization, methodology, editing manuscript, analysis – JH. Hardware provision, funding acquisition, supervision, conceptualization, methodology, analysis, editing manuscript – TC. Methodology, investigation, analysis, writing and editing manuscript – EOS. Investigation, analysis, writing manuscript – CC. Investigation, analysis – OT, PD, DH, KS. Hardware, experimental methodology – NH. Hardware, experimental methodology, analysis −PR. Data analysis −DS. Database resource – KC. Experimental methodology, analysis – DF,CB

BC,EOS,CC,JOS,OT,AS, KS, DF,PR,NH,DS,CB,TC,JH,LD report no conflict of interests to declare

