## Supplemental Figures and Tables 1-3 for "Kidney single-cell atlas reveals myeloid heterogeneity in progression and regression of kidney disease": merged_supplemental.pdf

### Supp fig 1

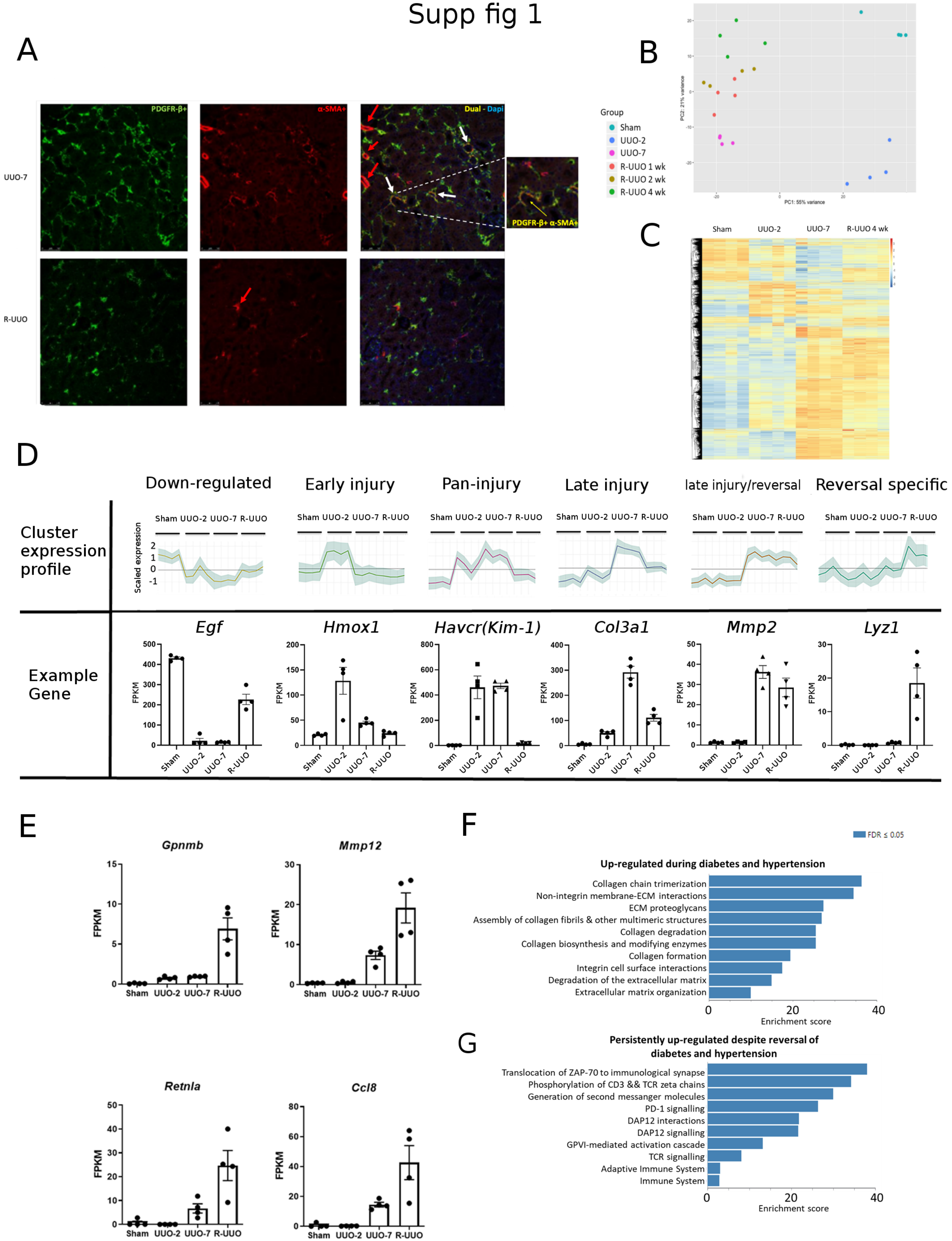
A
B
C
D
E
F
G
UUO-7
R-UUO
PDGFR-β+
α-SMA+
Dual - Dapi
PDGFR-β+ α-SMA+
Group
Sham
UUO-2
UUO-7
R-UUO 1 wk
R-UUO 2 wk
R-UUO 4 wk
PC1: 55% variance
PC2: 21% variance
Sham UUO-2 UUO-7 R-UUO 4 wk
Down-regulated
Early injury
Pan-injury
Late injury
late injury/reversal
Reversal specific
Cluster expression profile
Scaled expression
Sham UUO-2 UUO-7 R-UUO
Example Gene
Egf
Hmox1
Havcr(Kim-1)
Col3a1
Mmp2
Lyz1
FPKM
Gpnmb
Mmp12
Retnla
Ccl8
FDR ≤ 0.05
Up-regulated during diabetes and hypertension
Collagen chain trimerization
Non-integrin membrane-ECM interactions
ECM proteoglycans
Assembly of collagen fibrils & other multimeric structures
Collagen degradation
Collagen biosynthesis and modifying enzymes
Collagen formation
Integrin cell surface interactions
Degradation of the extracellular matrix
Extracellular matrix organization
Enrichment score
Persistently up-regulated despite reversal of diabetes and hypertension
Translocation of ZAP-70 to immunological synapse
Phosphorylation of CD3 && TCR zeta chains
Generation of second messanger molecules
PD-1 signalling
DAP12 interactions
DAP12 signalling
GPVI-mediated activation cascade
TCR signalling
Adaptive Immune System
Immune System
Enrichment score

Supp fig 2

A

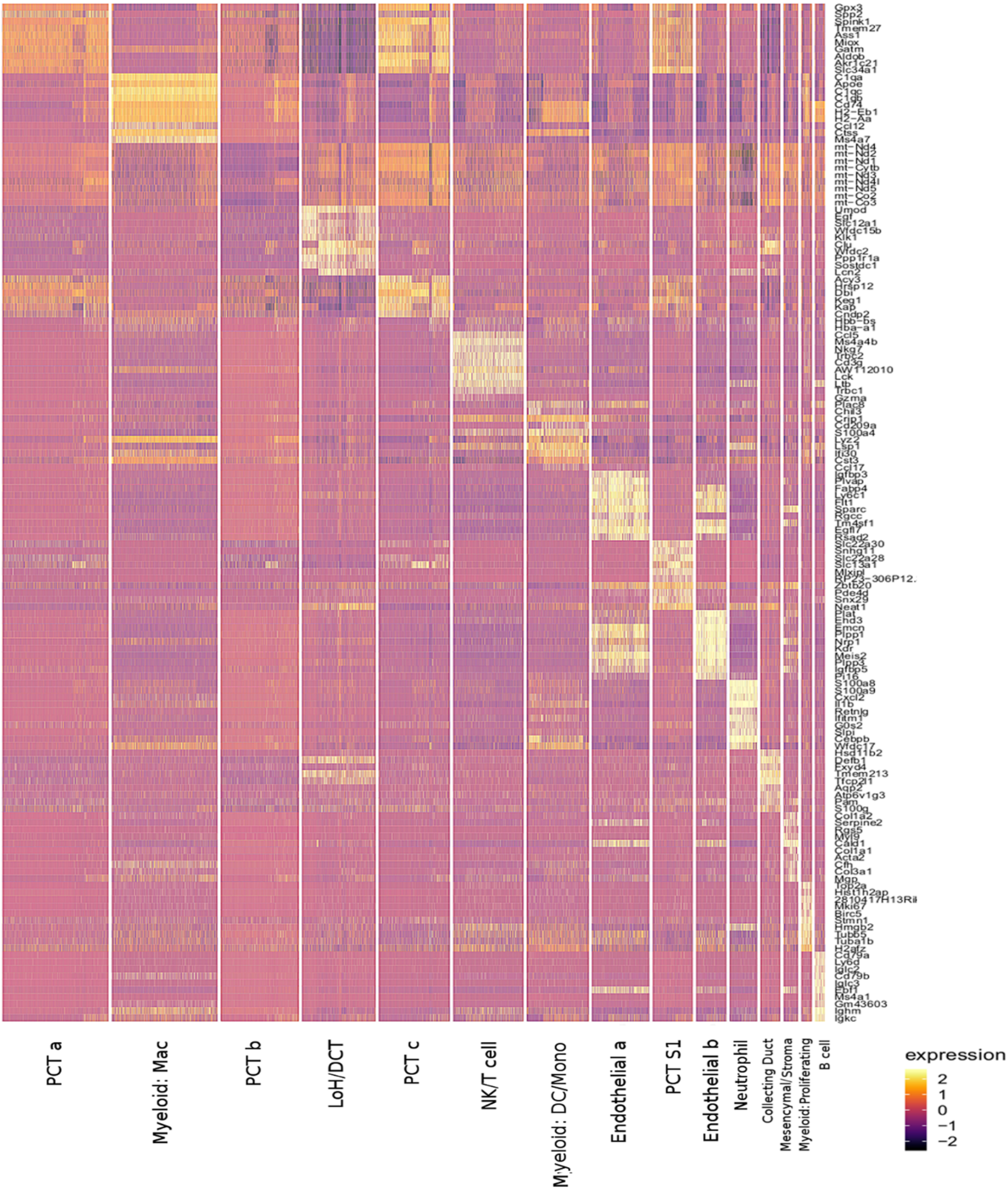
PCT a
Myeloid: Mac
PCT b
LoH/DCT
PCT c
NK/T cell
Myeloid: DC/Mono
Endothelial a
PCT S1
Endothelial b
Neutrophil
Collecting Duct
Mesencymal/Stroma
Myeloid:Proliferating
B cell
expression
2
1
0
−1
−2

B

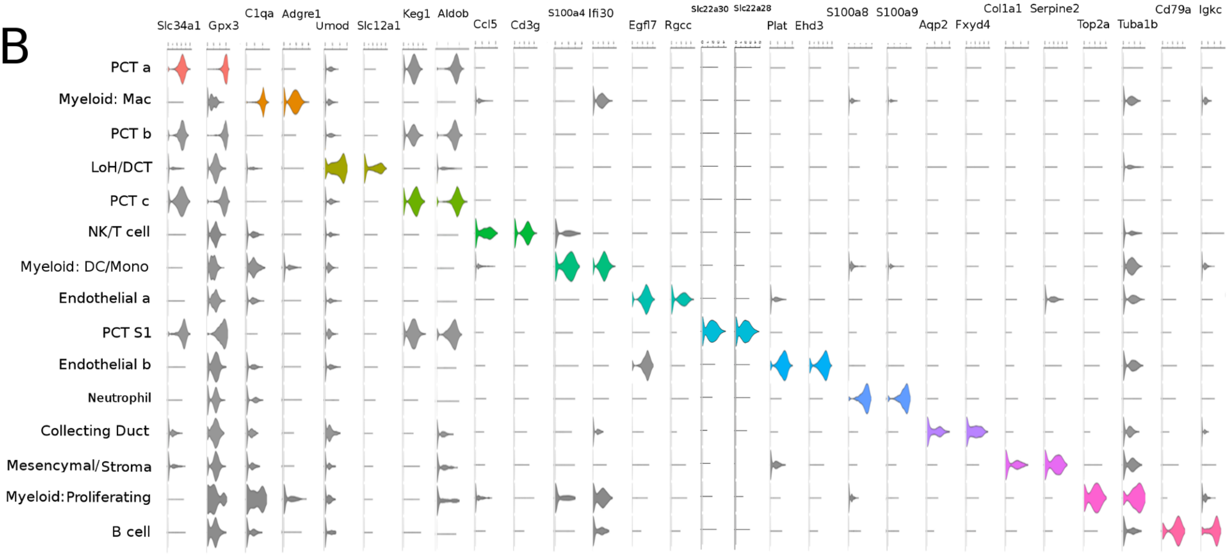
Slc34a1
Gpx3
C1qa
Adgre1
Umod
Slc12a1
Keg1
Aldob
Ccl5
Cd3g
S100a4
Ifi30
Egfl7
Rgcc
Slc22a30
Slc22a28
Plat
Ehd3
S100a8
S100a9
Aqp2
Fxyd4
Col1a1
Serpine2
Top2a
Tuba1b
Cd79a
Igkc
PCT a
Myeloid: Mac
PCT b
LoH/DCT
PCT c
NK/T cell
Myeloid: DC/Mono
Endothelial a
PCT S1
Endothelial b
Neutrophil
Collecting Duct
Mesencymal/Stroma
Myeloid:Proliferating
B cell

C

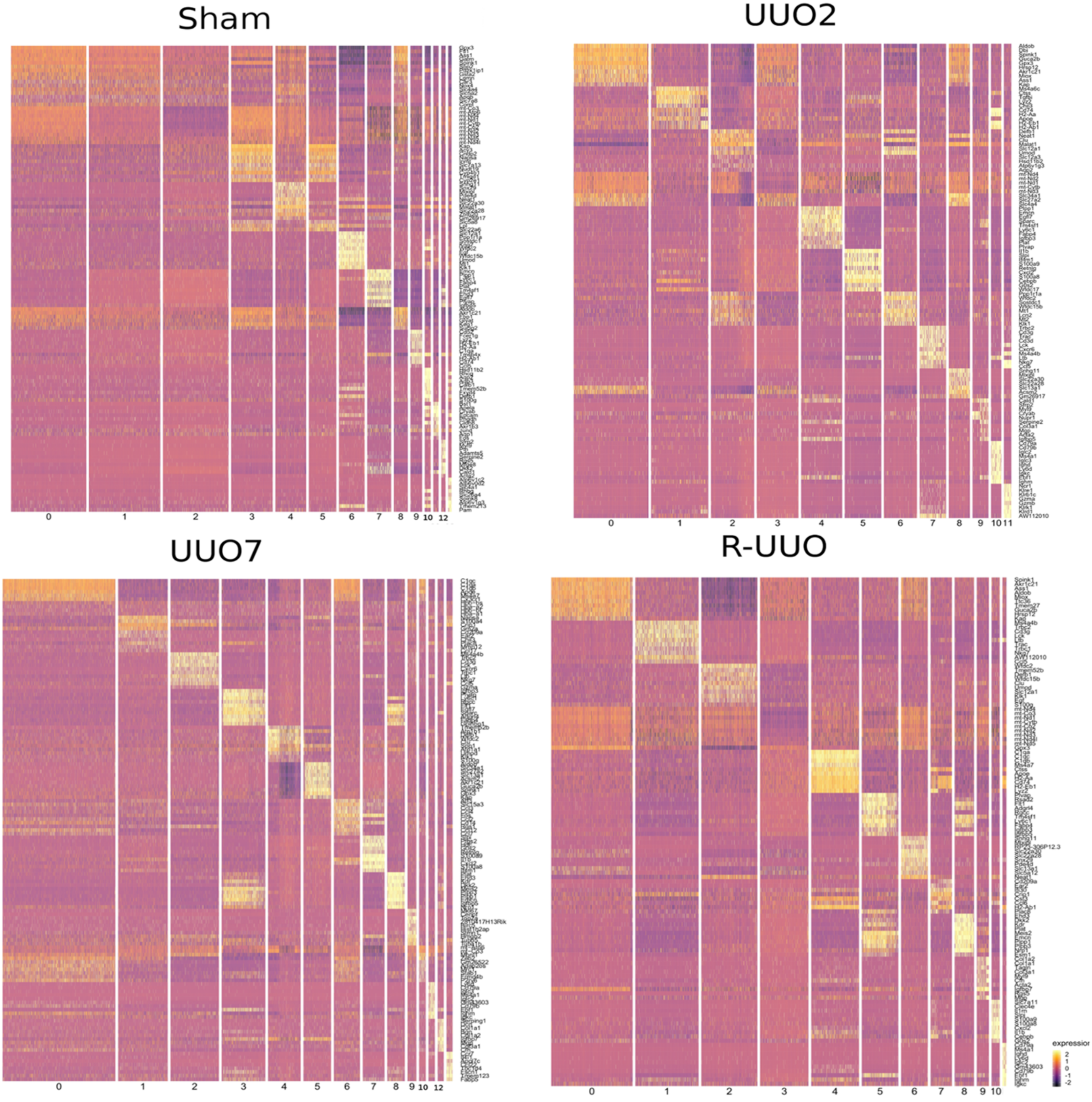
Sham
UUO2
UUO7
R-UUO
expression

D

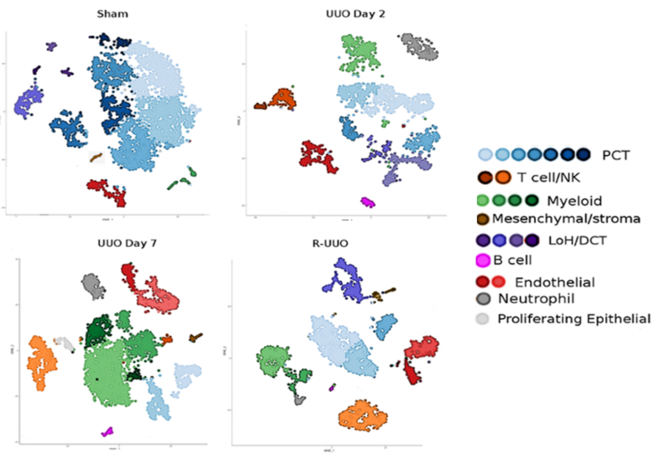
Sham
UUO Day 2
UUO Day 7
R-UUO
PCT
T cell/NK
Myeloid
Mesenchymal/stroma
LoH/DCT
B cell
Endothelial
Neutrophil
Proliferating Epithelial

E

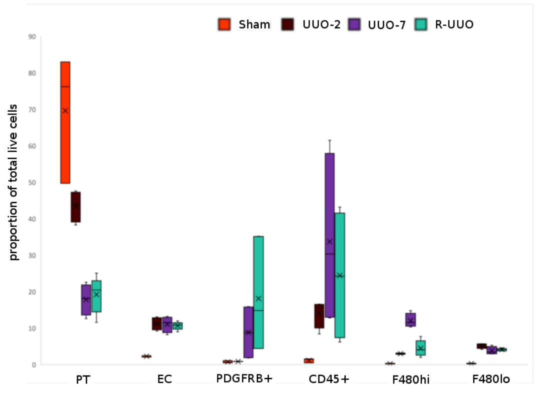
Sham
UUO-2
UUO-7
R-UUO
proportion of total live cells
PT
EC
PDGFRB+
CD45+
F480hi
F480lo

Supp fig 3

A

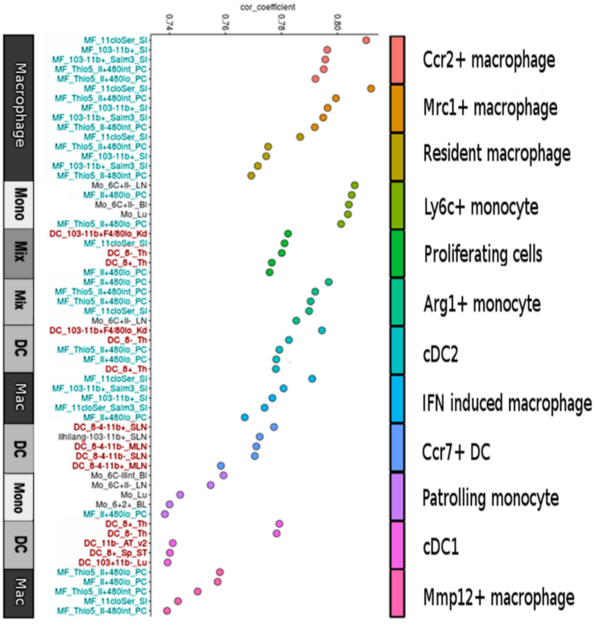
cor_coefficient
Macrophage
Mono
Mix
Mix
DC
Mac
DC
Mono
DC
Mac
Ccr2+ macrophage
Mrc1+ macrophage
Resident macrophage
Ly6c+ monocyte
Proliferating cells
Arg1+ monocyte
cDC2
IFN induced macrophage
Ccr7+ DC
Patrolling monocyte
cDC1
Mmp12+ macrophage

B

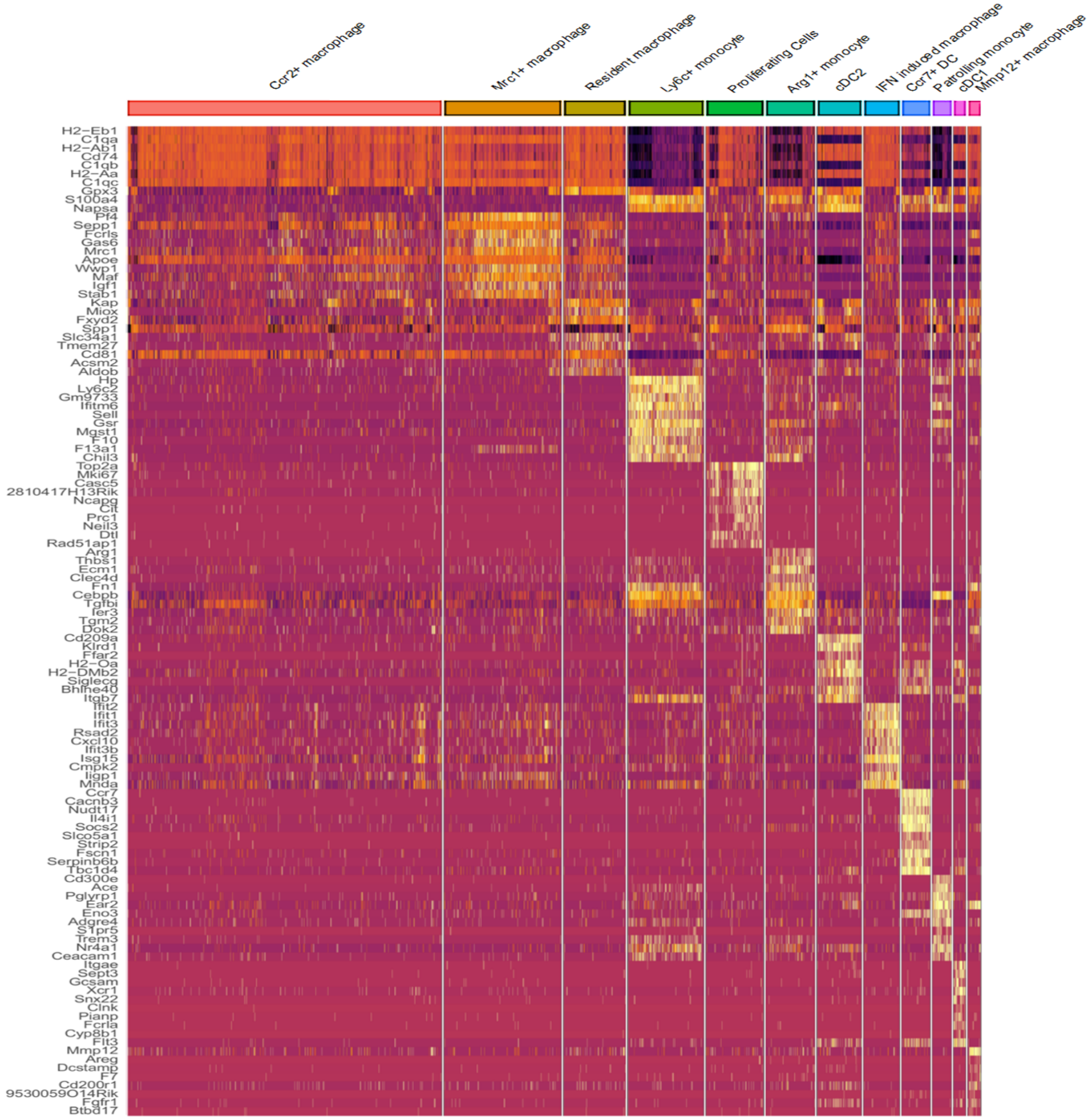
Ccr2+ macrophage
Mrc1+ macrophage
Resident macrophage
Ly6c+ monocyte
Proliferating Cells
Arg1+ monocyte
cDC2
IFN induced macrophage
Ccr7+ DC
Patrolling monocyte
cDC1
Mmp12+ macrophage
H2-Eb1
C1qa
H2-Ab1
Cd74
C1qb
H2-Aa
C1qc
Gpx3
S100a4
Napsa
Pf4
Sepp1
Fcrls
Gas6
Mrc1
Apoe
Wwp1
Maf
Igf1
Stab1
Kap
Miox
Fxyd2
Spp1
Slc34a1
Tmem27
Cd81
Acsm2
Aldob
Hp
Ly6c2
Gm9733
Ifitm6
Sell
Gsr
Mgst1
F10
F13a1
Chil3
Top2a
Mki67
Casc5
2810417H13Rik
Ncapg
Cit
Prc1
Neil3
Dtl
Rad51ap1
Arg1
Thbs1
Ecm1
Clec4d
Fn1
Cebpb
Tgfbi
Fer3
Tgm2
Dok2
Cd209a
Klrd1
Ffar2
H2-Oa
H2-DMb2
Siglecg
Bhlhe40
Itgb7
Ifit2
Ifit1
Ifit3
Rsad2
Cxcl10
Ifit3b
Isg15
Cmpk2
Iigp1
Mnda
Ccr7
Cacnb3
Nudt17
Il4i1
Socs2
Slco5a1
Strip2
Fscn1
Serpinb6b
Tbc1d4
Cd300e
Ace
Pglyrp1
Ear2
Eno3
Adgre4
S1pr5
Treml4
Nr4a1
Ceacam1
Itgae
Sept3
Gcsam
Xcr1
Snx22
Clnk
Pianp
Fcrla
Cyp8b1
Flt3
Mmp12
Areg
Dcstamp
F7
Cd200r1
9530059O14Rik
Fgfr1
Btbd17

C

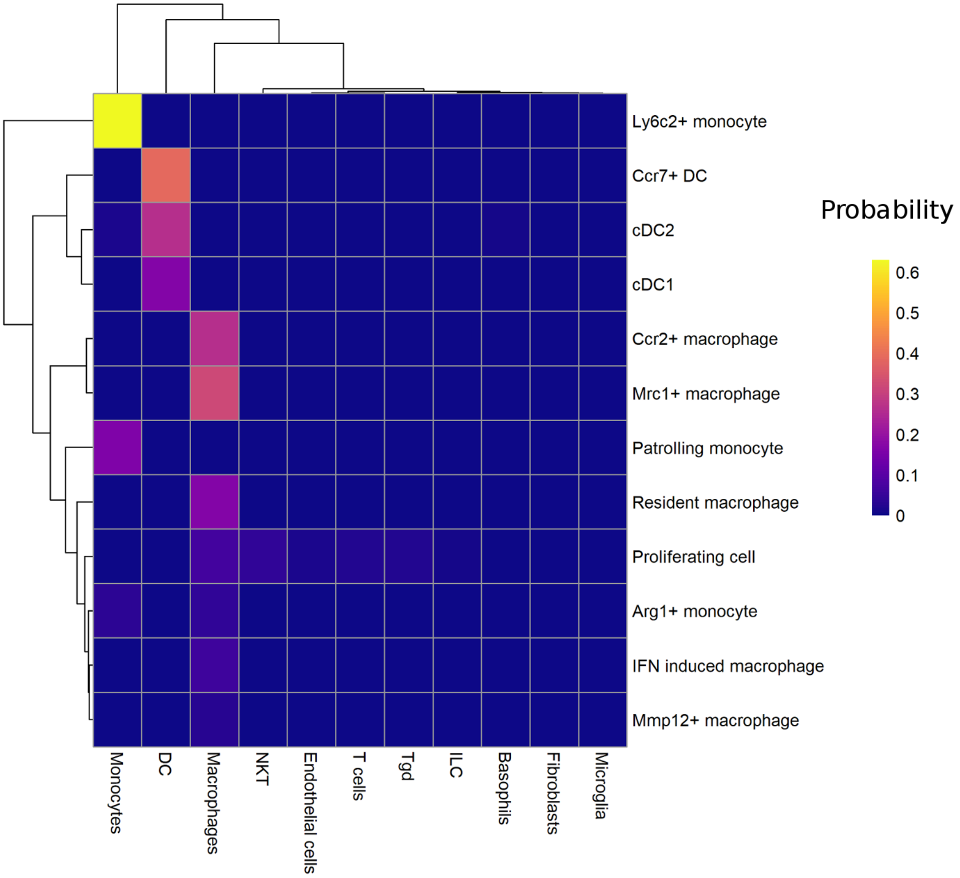

Ly6c2+ monocyte
Ccr7+ DC
cDC2
cDC1
Ccr2+ macrophage
Mrc1+ macrophage
Patrolling monocyte
Resident macrophage
Proliferating cell
Arg1+ monocyte
IFN induced macrophage
Mmp12+ macrophage
Monocytes
DC
Macrophages
NKT
Endothelial cells
T cells
Tgd
ILC
Basophils
Fibroblasts
Microglia
Probability
0.6
0.5
0.4
0.3
0.2
0.1
0

Supp fig 4

A

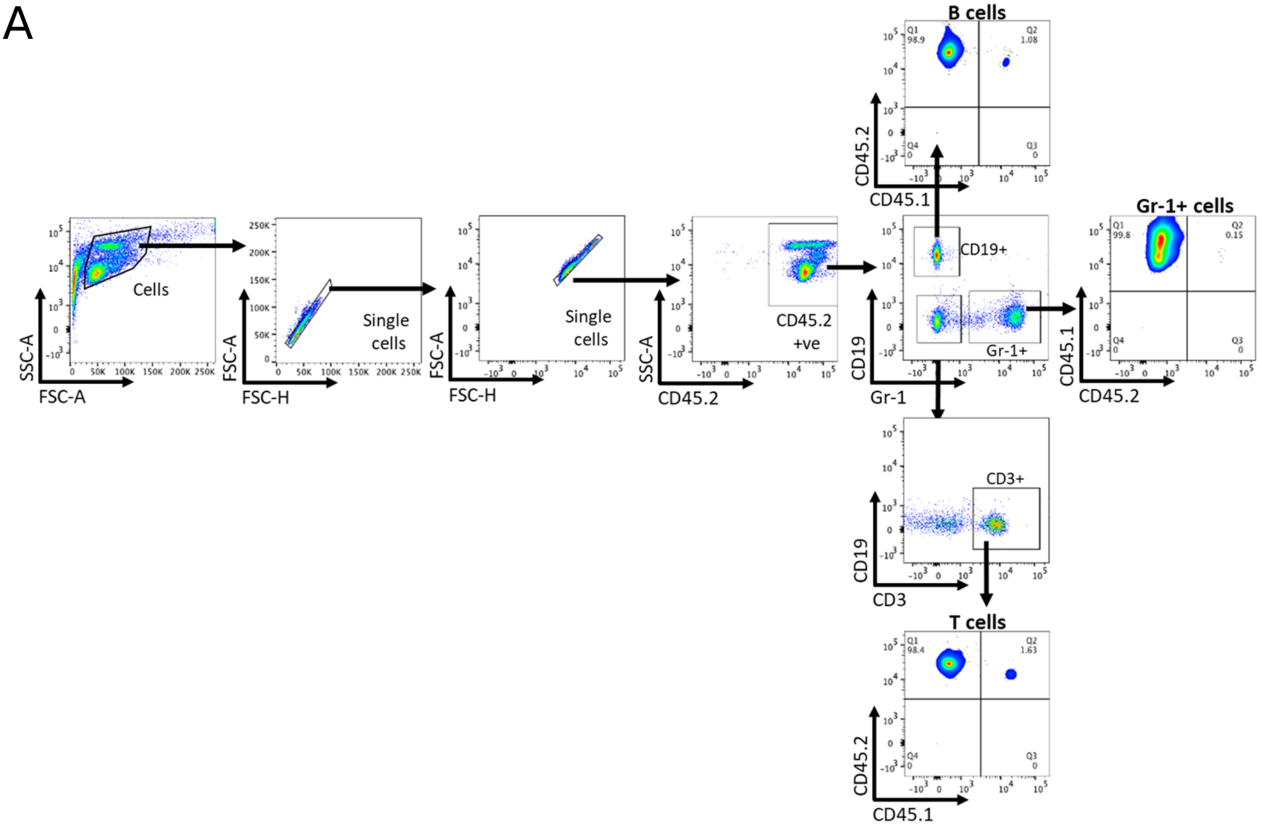

Cells
Single cells
Single cells
CD45.2 +ve
CD19+
Gr-1+
B cells
Gr-1+ cells
CD3+
T cells
SSC-A
FSC-A
FSC-A
FSC-H
SSC-A
CD45.2
CD19
Gr-1
CD45.2
CD45.1
CD45.1
CD45.2
CD19
CD3
CD45.2
CD45.1


B

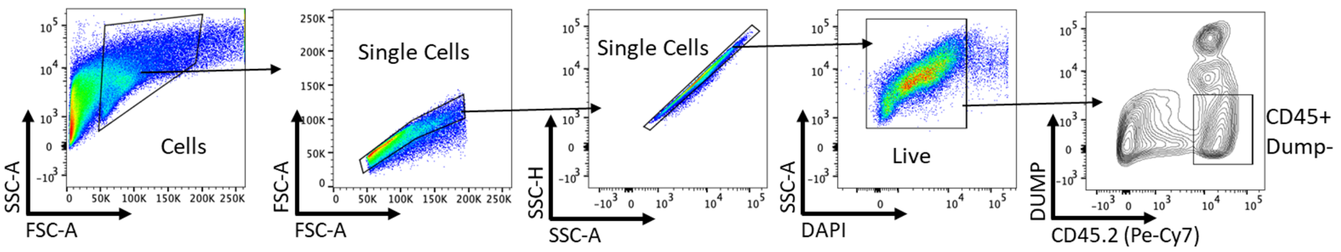

Cells
Single Cells
Single Cells
Live
CD45+ Dump-
SSC-A
FSC-A
FSC-A
SSC-H
SSC-A
SSC-A
DAPI
DUMP
CD45.2 (Pe-Cy7)

| Antibody | Company/Cat Num/Dilution |
|---|---|
| **Collagen III** | Southern Biotech/1330-08/1:200 |
| **PDGFR-β** | Abcam/ab32570/1:500 |
| **α-SMA** | Abcam/ab5694/1:400 |
| **F4/80** | Abcam/ab16911/1:100 |
| **YM1 (Chil3)** | Stemcell/60130/1:500 |
| **Mannose Receptor** | R&D/AF2535/1:200 |
| **Mmp12** | Santa Cruz/sc390863/1:100 |
| **Anti-rabbit biotinylated** | Vector Labs/BA1000/1:300 |
| **Anti-goat biotinylated** | Vector Labs/BA5000/1:300 |
| **Anti-rat biotinylated** | Vector Labs/BA9400/1:300 |
| **Anti-rabbit AF555** | ThermoFisher/A-11034/1:200 |
| **Anti-goat AF488** | ThermoFisher/A-11055/1:200 |
| **Anti-mouse AF488** | ThermoFisher/A-11029/1:200 |
| **Anti-rat AF488** | ThermoFisher/A-21434/1:200 |

Table 1: Antibodies utilised in immunohistochemistry/immunofluorescence studies.

| Gene | Probe/Assay ID |
|---|---|
| ***Mmp12*** | Mm00500554_m1 |
| ***Gpnmb*** | Mm01328587_m1 |
| ***Lyz1*** | Mm00657323_m1 |
| ***Hprt*** | Mm00446968_m1 |

**Table 2.** Taqman probes used for qRT-PCR in studies.

| Antibody | Clone/Flurochrome/final concentration |
|---|---|
| **Live** | DAPI / 1:1000 |
| **CD45** | 30-F11/BV650 or APC/1:100 |
| **CD45.1** | A20/FITC/1:200 K or 1:100 blood |
| **CD45.2** | 104/Pe-Cy7/1:200 K or 1:100 blood |
| **F4/80** | BM8/Pe-Cy7 or APC / 1:100 or 1:200 K or 1:100 blood |
| **MHCII** | M5-114.15.2/APC-Cy7 / 1:400 |
| **Ly6G** | 1A8/BV421 / 1:200 |
| **Ly6C** | HK1.4/AF700 / 1:200 |
| **CD11b** | M1_70/PE Dazzle / 1:1000 |
| **CD11c** | N418/BV605 / 1:100 |
| **CD64** | X54-5/7.1/PerCp5.5/1:200 |
| **CD24** | 30-F1/PE/1:200 |
| **Siglec-F** | E50-2440/BV421/1:200 |
| **TCRβ** | H57-597/BV421/1:200 |
| **CD19** | 6D5/BV421/1:200 |
| **CD3** | 17A2/PerCp5.5 1:100 |
| **CD206** | MR5D3/APC / 1:200 |
| **GR-1** | RB6-8C5/PE/1:200 |
| **CCR2** | SA203G11/BV605/1:200 |
| **PDGFRβ** | APB5/PE/1:50 |
| **CD31** | 390/BV605/1:100 |
| **LTL** | FL-1321/FITC/1:200 |

**Suppl Table 3**. Antibodies utilised in Flow Cytometry.

**Supplemental figure 1**

**a.** Immunofluorescence for PDGFR-$\beta^+$ and $\alpha$-SMA$^+$ indicates that a proportion of PDGFR-$\beta^+$ cells co-express $\alpha$-SMA$^+$ at UUO-7 (white arrows) indicating activation to myofibroblasts; 2 weeks following R-UUO, the interstitial PDGFR-$\beta^+$ cells no longer express $\alpha$-SMA$^+$, which is now restricted to the arterial smooth muscle walls (red arrows). **b.** Principal component (PC) analysis of RNA sequencing data from bulk cortical kidney tissue from animals undergoing sham surgery, or at UUO-2, UUO-7, or 1- 2- or 4-weeks post R-UUO (n=4/time-point). **c.** Heatmap of differentially expressed genes created using Pheatmap R package. Rows were clustered via unsupervised hierarchical clustering. **d.** Expression of exemplar genes aligning to each cluster **e.** Expression of *Gpnmb*, *Mmp12*, *Retnla* and *Ccl8* genes in RNAseq dataset **f.** Pathways enriched for genes that are induced in the kidney of *Cyp1a1mRen2* rats during diabetes and hypertension, but which revert towards baseline following tight glycaemic and blood pressure control (32) **g.** Pathways enriched for genes that are persistently elevated in the kidney of *Cyp1a1mRen2* rats despite onset of tight glycaemic and blood pressure control

**Supplemental figure 2**

**a** Top 10 differentially expressed genes by fold change in each cluster, calculated using Wilcoxon signed-rank test. The colour scheme is based on z-score distribution. **b** Violin plots showing the expression levels of selected marker genes in each cluster. The x axis shows the log-scale normalized read count. **c** Top 10 differentially expressed genes by fold change in each cluster across the sham, UUO-2, UUO-7 and R-UUO libraries analysed individually using Wilcoxon signed-rank test. The colour scheme is based on z-score distribution **d** Separate tSNE plots restricted to cells from each separate time-point coloured by shared nearest neighbour (SNN) allocated cluster, annotated by cell type **e** Flow cytometry was employed to determine the proportion of total kidney cells at each time point comprised by proximal tubular cells (PT), endothelial cells (EC), fibroblasts (PDGFR-β), immune cells (CD45), and F4/80hi or F4/80lo macrophages.

**Supplemental figure 3**

**a** Top 5 Immunological Genome Project (ImmGen) reference cell types per cluster ranked by Spearman's correlation coefficients with full dataset details following a cluster-to-references analysis using cluster identify predictor v2. **b** Top 10 differentially expressed genes by fold change per cluster, calculated using Wilcoxon signed-rank test. The colour scheme is based on z-score distribution **c** Consensus matrix of ImmGen reference cell types using SingleR generated labels (bottom) with final classifications

**Supplemental figure 4**

**a** Flow cytometry gating strategy for characterizing circulating blood immune cells into B-cells ($CD19^+$), monocyte/neutrophils ($Gr\text{-}1^+$), and T-cells ($CD3^+$) using the D2 blood sample as an exemplar. **b** Flow cytometry gating strategy for sorting myeloid cells in the kidney. Dump gate includes TCRβ, CD19, Ly6G and SiglecF all on the same fluorophore to exclude T-cells, B-cells, neutrophils and eosinophils respectively.

**Supplemental Table 1**

Antibodies utilised in immunohistochemistry/immunofluorescence studies.

**Supplemental Table 2**

Taqman probes used for qRT-PCR in studies.

**Supplemental Table 3**

Antibodies utilised in Flow Cytometry

**Supplemental Table 4**

Summary of 10x sequencing statistics

**Supplemental Table 5**

Bulk RNA-Seq matrix of differentially expressed genes in each gene cluster,
expressed in FKPM

**Supplemental Table 6**

Single cell RNA-Seq table of differentially expressed genes in the global data as described in figure two, expressed per cluster, where avg_logFC is the average log fold change between the cluster in question and the remaining clusters, pct.1 and pct.2 is the percentage of cells expressing the gene in question in the cluster in question the remaining clusters, and p_val_adj is the adjusted P-value following Bonferroni correction for multiple hypothesis testing.

**Supplemental Table 7**

Single cell RNA-Seq table of differentially expressed genes in the myeloid only data as described in figure three, expressed per cluster, where avg_logFC is the average log fold change between the cluster in question and the remaining clusters, pct.1 and pct.2 is the percentage of cells expressing the gene in question in the cluster in question the remaining clusters, and p_val_adj is the adjusted P-value following Bonferroni correction for multiple hypothesis testing and rank is the assigned rank for gene set enrichment analysis calculation.
